# Aquatic biodiversity enhances multiple nutritional benefits to humans

**DOI:** 10.1101/691444

**Authors:** Joey R. Bernhardt, Mary I. O’Connor

## Abstract

Humanity depends on biodiversity for health, well-being and a stable environment. As biodiversity change accelerates, we are still discovering the full range of consequences for human health and well-being. Here, we test the hypothesis -- derived from biodiversity - ecosystem functioning theory -- that species richness and ecological functional diversity allow seafood diets to fulfill multiple nutritional requirements, a condition necessary for human health. We analyzed a newly synthesized dataset of 7245 observations of nutrient and contaminant concentrations in 801 aquatic animal taxa, and found that species with different ecological traits have distinct and complementary micronutrient profiles, but little difference in protein content. The same complementarity mechanisms that generate positive biodiversity effects on ecosystem functioning in terrestrial ecosystems also operate in seafood assemblages, allowing more diverse diets to yield increased nutritional benefits independent of total biomass consumed. Notably, nutritional metrics that capture multiple micronutrients essential for human well-being depend more strongly on biodiversity than common ecological measures of function such as productivity, typically reported for grasslands and forests. Further, we found that increasing species richness did not increase the amount of protein in seafood diets, and also increased concentrations of toxic metal contaminants in the diet. Seafood-derived micronutrients are important for human health and are a pillar of global food and nutrition security. By drawing upon biodiversity-ecosystem functioning theory, we demonstrate that ecological concepts of biodiversity can deepen our understanding of nature’s benefits to people and unite sustainability goals for biodiversity and human well-being.

**Significance statement:** Food security is not simply about maintaining yields, it is also about the need for a stable supply of nutritionally diverse foods. Obtaining nutritious food is a major challenge facing humanity and diverse aquatic ecosystems can help meet this goal. To test how aquatic biodiversity affects human health, we assembled a new dataset of nutrients, contaminants and ecological traits of 801 aquatic species. We used ecological models to quantify the role of species richness and ecological functional diversity and found that these biodiversity dimensions enhanced seafood micronutrient provisioning by the same mechanisms that link biodiversity to productivity in grasslands, forests and other systems. Our results underscore the need to minimize aquatic biodiversity loss to sustain and improve human well-being.

## Introduction

Species losses and range shifts due to climate change, harvesting, and other human activities are altering aquatic biodiversity locally and globally (1–5). In aquatic ecosystems, not only are some species severely depleted because of overfishing or habitat loss (3, 6–8), the ecosystem-level dimensions of biodiversity such as the total number of species and their functional diversity have also changed (9). Beyond the loss of particular species, changes in ecosystem-level dimensions of biodiversity threaten numerous ecosystem services to humans, which include the cultural, economic or health benefits people derive from nature (10–13). In many regions, such as tropical coastal systems, the cumulative impacts of human activities are severe and associated with strong declines in taxonomic and ecological functional diversity (6) and coincide with regions with high dependence of people upon wild-caught seafood for food and nutrition (14). In temperate regions, where some coastal communities depend on local wild seafood harvests to meet their nutritional needs (15, 16), species richness may be increasing as species recover from exploitation and warmer oceans allow species to expand their ranges into new territory (1, 2, 17).

There is growing concern that biodiversity change leads to changes in human health and well-being (10, 13, 18). Specific and quantitative links between aquatic biodiversity and human health that distinguish contributions of species diversity from those of biomass, as predicted by biodiversity-ecosystem functioning theory, have not been established. At a time of unprecedented global change and increasing reliance on seafood to meet nutritional demands (19), there is an urgent need to understand how changing aquatic ecosystem structure may alter the provisioning of seafood-derived human nutrition.

Seafood, consisting of wild-caught marine and freshwater finfish and invertebrates, provides an important source of protein and calories to humans. Additionally, unlike staple foods such as rice or other grains, seafood can address multiple dimensions of food and nutritional security simultaneously by providing essential micronutrients, such as vitamins, minerals and polyunsaturated essential fatty acids critical to human health (19–22). Given the multiple attributes of seafood valuable to human health, it is possible that the diversity of an aquatic assemblage, distinct from the inclusion of any particularly nutritious species, could support human well-being consistent with a large body of evidence for biodiversity’s major contributions to ecological functions (11, 23–26). Dietary diversity is a basic tenet of a nutritious diet (27) and it is widely appreciated that diets composed of more food groups and more species are more nutritious (28–31). Ecological measures of dietary diversity (diet diversity, species richness, functional diversity and Simpson’s index of evenness) have been associated with nutritional value of diets in a range of contexts (27, 29, 32–38). These studies rely on relationships between species included in the diet (or other food intake measures) and nutritional adequacy of reported diets. However, a simple correlation between dietary diversity and a measure of dietary benefits provides only partial support for a claim that biodiversity benefits human well-being consistent with the same ecological processes by which biodiversity supports numerous ecosystem functions and services (23, 26). We build upon this foundation of empirical relationships between diet diversity and diet quality by placing this question in the quantitative ecological theoretical framework that relates biodiversity to function (24, 25), thereby laying the groundwork for additional development of links between biodiversity science and our understanding of human well-being.

Ecological theory predicts that biodiversity can be ecologically and economically important, apart from the importance of total biomass or the presence of particular species (23, 39). According to theory and over 500 explicit experimental tests (23, 40, 41) diversity in ecological communities and agricultural systems enhances ecosystem functioning by two mechanisms: (i) more diverse assemblages may outperform less diverse assemblages *of the same density or biomass* of individuals because more diverse assemblages will include more of the possible species and are therefore more likely to include high-performing species, assuming random processes of including species from the species pool (a *selection effect*), or (ii) more diverse assemblages of a given density (or biomass) contain species with complementary functional traits, allowing them to function more efficiently (a *complementarity effect*) (25, 39). For aquatic animals, increased diversity enhances productivity of fish biomass (42) and also enhances temporal stability of biomass production and total yields (43, 44), providing economic and nutritional benefits to humans related to increased stability of harvests and production of biomass for consumption (43). However, when considering aquatic species from the perspective of human nutrition, functions other than biomass production become relevant because total seafood biomass consumption is not predictive of micronutrient benefits from seafood (45, 46).

Here we test a hypothesis central to ecological theory in the 21st century: whether biodiversity *per se* (species richness and ecological functional diversity), distinct from the identities and abundance of species, enhances human well-being (**Figure 1**). We chose a measure of human well-being distinct from provision of protein, calories or total yields - the micronutrient benefits of seafood. For increasing biodiversity *per se* (as opposed to increasing total seafood consumption) to enhance nutritional benefits as predicted by biodiversity-ecosystem functioning theory (25, 47), the amounts of various micronutrients within edible tissues must differ among species and, furthermore, micronutrient concentrations must trade off among species, such that species that have relatively high concentrations of some nutrients also have relatively low concentrations of others (25). Specifically, a ‘biodiversity effect’ (*sensu* (*25*)) on nutritional benefits requires that concentrations of multiple nutrients are negatively correlated with each other, or uncorrelated, when compared among species, creating a complementary distribution of micronutrients across species. In contrast, if micronutrient concentrations in edible tissue are positively correlated for multiple micronutrients across species such that, for example, a species containing high amounts of iron also has a high essential fatty acid concentration, thereby containing multiple nutrients in high concentrations simultaneously, seafood species or ecological functional *diversity* in the diet would not be important. In the case of positive correlations among micronutrient concentrations, the ecosystem service of nutritional benefits would be enhanced by consuming more fish biomass or by selecting a few highly nutritious species, without considering species richness or ecological functional diversity.

**Figure 1.**
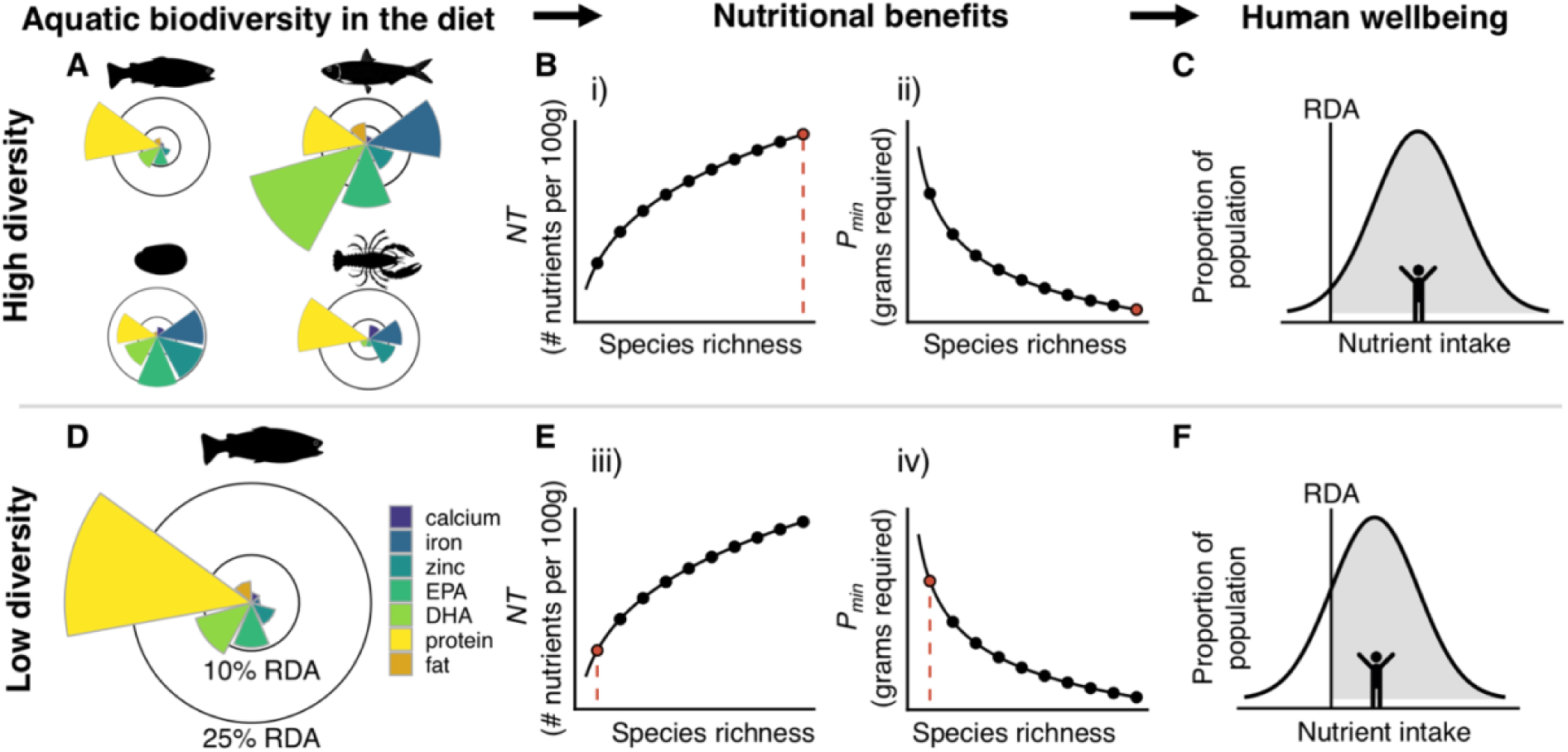
Aquatic biodiversity increases human well-being because edible species have distinct and complementary multi-nutrient profiles (**A**) and differ in mean micro- and macronutrient content (shown here relative to 10% and 25% thresholds of recommended dietary allowance (RDA) guidelines) for representative finfish (*Abramis brama*, *Mullus surmuletus*), mollusc (*Mytilus galloprovincialis*) and crustacean species (*Nephrops norvegicus*). Biodiversity – ecosystem functioning theory predicts that nutritional benefits, including the number of nutrient RDA targets met per 100g portion (*NT*; i, iii) and minimum portion size (*P_min_*; ii, iv) (**B, E**), are enhanced with increasing seafood species richness. Orange dots in panels B and E correspond to potential diets high and low biodiversity levels. Seafood consumers with limited access to seafood each day may not reach RDA targets if diets are low in diversity (**F** vs **C**; grey shading indicates proportion of population that meets nutrient requirements). DHA: docosahexaenoic acid, EPA: eicosapentaenoic acid.

We aimed to bridge two distinct theoretical frameworks - the biodiversity-ecosystem functioning theory and human nutrition science - by quantitatively testing for effects of aquatic species richness and ecological functional diversity (48, 49) in seafood diets on nutritional benefits via complementarity or selection effects. We used the public health measure of recommended dietary allowance (RDA) index to quantify nutritional benefits. RDAs are nutrient-based reference values that indicate the average daily dietary intake level that is sufficient to meet the nutrient requirement of nearly all (97 to 98 percent) healthy individuals in a particular life stage and gender group (50). Here we used the RDA for females aged 19-50 (SI Appendix, Tables S1, S2). We measured nutritional value in terms of concentrations relative to RDAs, and we refer to these recommended amounts (or portions thereof) as ‘RDA targets’ (SI Appendix, Table S1, Table S2, Methods 1.1). We quantified nutritional value in two ways: 1) the minimum amount of seafood tissue (in grams, g) required to meet given RDA targets (either for a single nutrient, or five micronutrients simultaneously; referred to as ‘minimum portion size required’, *P_min_* (SI Appendix, Table S1, Equation A1, Methods 1.1) and 2) the number of nutrients that meet an RDA target in a single 100g seafood portion (*NT*, Table S1, Equation A2). By considering nutritional value per unit biomass in both metrics, we avoided confounding diversity of seafood consumed with the total amount consumed (Methods 1.3). We first tested two hypotheses: (i) seafood species richness increases *NT* due to complementarity in nutrient concentrations among species and (ii) seafood species richness increases the nutritional value of a 100g edible portion of seafood, thereby lowering the minimum portion size, *P_min_*, and improving the efficiency with which seafood consumers reach nutritional targets (**Figure 1**). Following biodiversity-ecosystem functioning theory, we predicted that increased species richness is correlated with ecological functional diversity (51) in potential seafood diets, and that ecological functional diversity is related to diversity in the concentration of essential elements and fatty acids that have nutritional value to human consumers, such that species and ecological functional diversity yields increased nutritional benefits. We also tested the hypothesis (iii) that seafood diversity increases total intake of heavy metal contaminants, because some aquatic animals are known to bioaccumulate toxic metals in their tissues. For this reason, variation in bioaccumulation among species could lead to a biodiversity effect on contaminant intake that is detrimental to human health.

In a global analysis of over 5040 observations of nutrient concentrations in 547 aquatic species (SI Appendix, **Figure S1**), we considered provision of nutritional benefits to human consumers. To assess whether the relationships between biodiversity and human nutrition benefits depend on the geographic extent (global, local) over which seafood are harvested or accessed (11), we tested whether seafood species richness is associated with higher nutritional value at local scales (*vs* global scale) in traditional indigenous seafood diets in North America (SI Appendix, Methods 1.4). Seafood is critical for indigenous groups, who on average consume seafood at a rate that is 15 times higher than the global average *per capita* consumption rate (16). To test our hypotheses at the geographic scale of local consumer communities, we complemented our global analysis with additional analyses of 25 – 57 species in fourteen geographically constrained groups of species consumed together as part of traditional indigenous diets (SI Appendix, Methods 1.4).

## Results

### Diversity of seafood nutrient concentrations

Biodiversity effects via complementarity or selection require that species differ in their functional traits. We found that the global species pool was highly diverse with regard to concentrations of the micronutrients iron, zinc, calcium, and two fatty acids, docosahexaenoic acid (DHA) and eicosapentaenoic acid (EPA), in edible fish tissue relative to RDAs for those micronutrients (**Figure 2**; micronutrient geometric coefficients of variation (CV): ln(iron) = 3.28, ln(calcium) = 3.56, ln(EPA) = 2.62, ln(zinc) = 3.02, ln(DHA) = 2.17; note the log scale). We observed limited variation in protein concentrations (ln(protein) CV = 0.04). The frequency distribution of trait values, such as nutrient concentrations, across species may indicate the potential strength of biodiversity effects, with lognormal distributions (such as we observed for micronutrients) more likely to confer strong effects of biodiversity than normal distributions with low dispersion (as we observed for protein). Most species did not meet a single micronutrient RDA in a 100g portion: fewer than half of the 547 species we examined reached an RDA target of 10% RDA for calcium, iron and the essential fatty acid EPA in a standard 100g edible portion of a single species (SI Appendix, Table S3).

**Figure 2.**
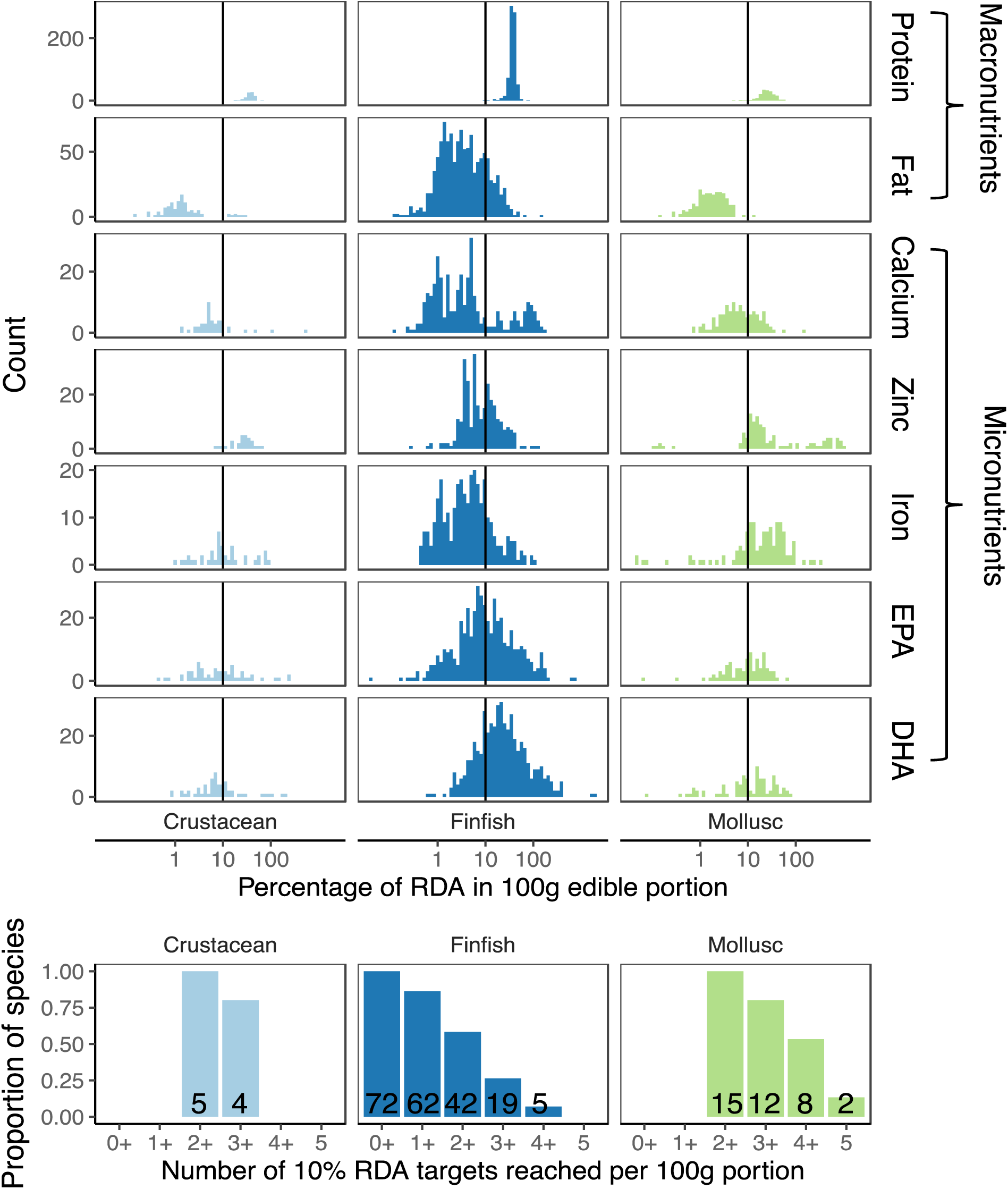
Variation in nutrient concentrations differs among taxonomic groups. (**A**) Frequency of reported protein, fat and micronutrient (including eicosapentaenoic acid (EPA), docosahexaenoic acid (DHA)) content in 100 g of the edible portion of 547 seafood species. Note the x-axis is plotted on a log scale. (**B**) Proportion of species, and number shown on each bar, with available data that reach 10% of RDA targets for any one, two or up to five of the micronutrients examined here.

### Biodiversity increased the nutritional content of an edible portion of seafood

We found that seafood species richness not only enhanced nutritional value for consumers selecting seafood from our global species dataset, but that seafood species richness *per se* was essential to meeting nutritional targets for seafood diets with limited biomass consumption. Increasing seafood species richness allowed simulated diets to reach more RDA targets per 100g of tissue, so that nutritional value increased with species richness even as total biomass consumption (e.g., total seafood portion size) remained constant. We quantified the minimum amount of seafood, in grams, that would be required to reach an RDA target (SI Appendix, Table S2) at each of ten levels of species richness (referred to as minimum portion size, *P_min_*, for which lower values signify higher nutrition benefits to consumers per gram seafood consumed, Methods 1.3). We then estimated the biodiversity effect using Equation 3 (Methods 2.3), in which *b_Pmin_* is the scaling coefficient that describes how function (here, *P_min_*) varies with species richness; higher absolute values of *b_f_* (where *f* is an ecosystem function) indicate a steeper relationship between biodiversity and function, and can be used to compare ‘biodiversity effects’ among studies and systems (12, 52). As species richness increased in potential diets, *P_min_* declined, and RDA targets for each micronutrient were achieved with less total seafood intake (**Figure 3A**, *b_Pmin_* < 0 for every micronutrient: calcium −0.32 (95% CI −0.35, −0.28), iron −0.24 (95% CI −0.27, −0.22), zinc −0.26 (95% CI −0.28, −0.23), EPA −0.25 (95% CI −0.27, −0.23) and DHA −0.22 (95% CI −0.23, −0.21)). Increasing species richness reduced the minimum portion size required, *P_min_*, in our sample diets, independent of systematic changes in the identity of species included (see Methods 2.2), because the diets were assembled using random samples of the species pool. The restricted variation in protein concentrations (**Figure 2A**), combined with high levels of protein in all edible tissues, lead to no benefit, and a minimal detrimental effect of seafood species richness on protein provisioning (**Figure 3A***, b_Pmin_* = 0.0071 95% CI 0.0062, 0.0080). In other words, the ecosystem service of protein provisioning was adequately provided by total seafood edible biomass, and not improved by species richness or even species identity. These findings are consistent with demonstrations that variety and diversity in diets is important for nutrition (28, 29), but extend these findings to show seafood species richness allows consumers to gain more nutritional benefit without consuming more total seafood biomass, and explicitly relates this pattern with general effects of biodiversity in ecological systems.

**Figure 3.**
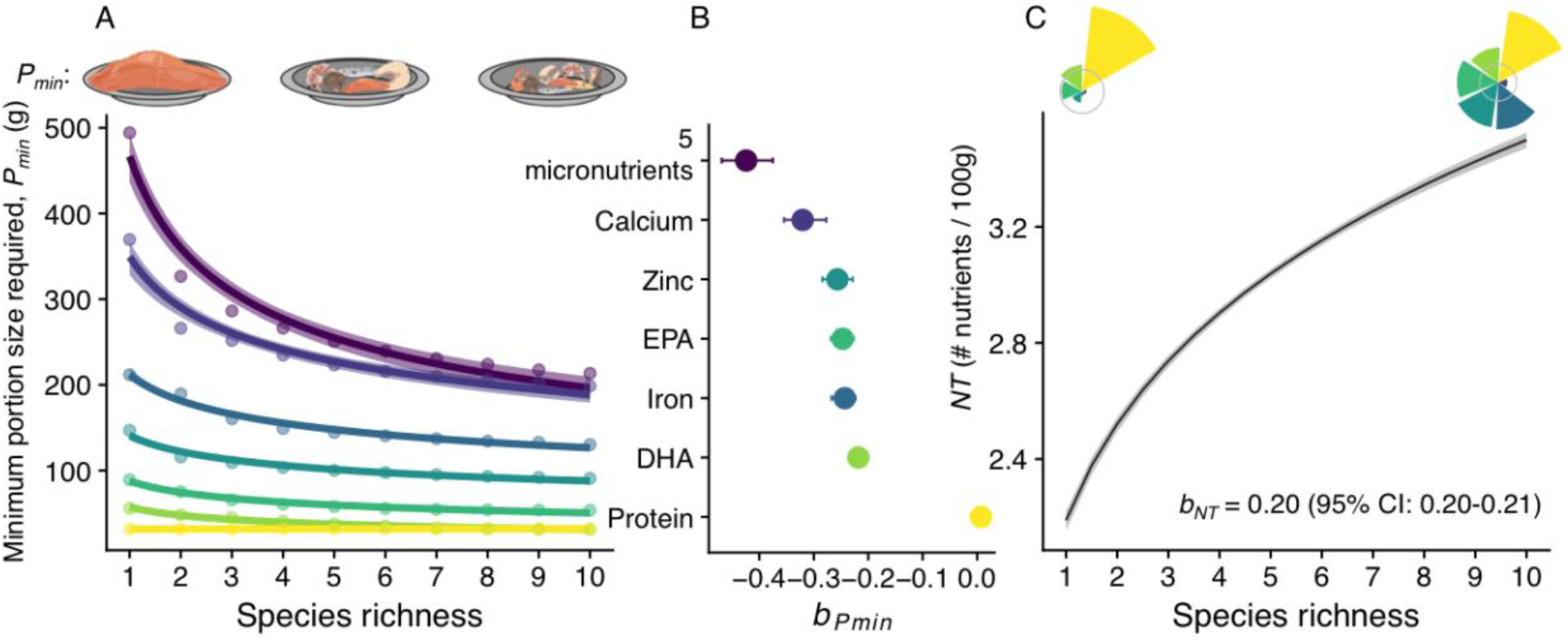
Aquatic biodiversity enhances nutritional benefits. (**A**) Seafood species richness improves the efficiency with which human diets can meet RDA targets by reducing the minimum portion size required, *P_min_*, to meet RDA targets (measured in grams of seafood). *P_min_* is shown for five micronutrients and protein separately (points are median values for calcium, iron, zinc, EPA, DHA and protein, lines show the fit of Equation 3 to the data and shading refers to 95% CI) as well as for five micronutrients simultaneously (top purple line). Colours corresponding to each nutrient in panel A are shown in panel B. (**B**) Estimates (± 95% CI) for the scaling parameter that relates species richness to *P_min_* (*b_Pmin_*) (Equation 3). (**C**) Species richness increases the number of distinct nutrient RDA targets met in a 100g seafood portion (*NT*); black line and 95% confidence interval correspond to the fit of Equation 2 to mean *NT* derived from resampled diets from the global seafood species pool. Flower plots in **C** summarize micronutrient concentrations relative to 10% RDA (grey circle) in two representative diets at low and high species richness levels. Data shown in panels A and C are derived from *n* = 1000 resampled diets.

We then considered the effects of seafood species richness on the provisioning of multiple nutrients simultaneously (Methods 1.3). This is referred to as a multifunctional benefit of biodiversity (53, 54), and takes into account possible trade-offs or correlations among functions, in this case, concentrations of micronutrients. For some ecosystem services (e.g. water quality or ecotourism), benefits of biodiversity accumulate when multiple ecosystem functions are considered simultaneously (54–56). Consistent with biodiversity-ecosystem functioning theory, we found that in the case of a multifunctional metric of an ecosystem service defined from the human beneficiary’s perspective (i.e. multiple micronutrient targets reached simultaneously), biodiversity benefits for the multifunctional service are greater than for individual functions (*b_Pmin_* for all five micronutrients simultaneously = −0.42 (95% CI −0.47, −0.38) vs. single nutrients *b_Pmin_* range from −0.32 (95% CI −0.35, −0.28) for calcium to −0.22 for DHA (95% CI −0.23, - 0.21)) that comprise the ecosystem service (**Figure 3A, B**). Increasing seafood species richness from one to ten species in 1000 simulated, resampled diets drawn from our global species pool allowed diets to meet RDA targets for five essential microelements and fatty acids simultaneously more than twice as efficiently (i.e. a median of 494.19 g of tissue required with one species vs. median of 213.34 g of tissue required with ten species) (**Figure 3A, B**). Then, we assessed the effects of biodiversity when the total biomass of seafood consumed was held constant, at 100g, by counting the number of nutrients for which RDA targets were reached in a 100g portion (*NT*). We found positive effects of biodiversity on the number of nutrients that met RDA targets, *NT*, in a single 100g portion (**Figure 3C**): diets with higher species richness reached more nutritional targets (higher *NT*) per 100g serving than diets of the same fish biomass comprising fewer species (*b_NT_* = 0.20 (95% CI: 0.20-0.21) **Figure 3C**). These findings were robust to different RDA target levels (for *P_min_*, they were independent of RDA target level, and for *NT* they were positive over the range from 1% RDA to 40% RDA per 100g portion, SI Appendix, **Figure S2**). These results reveal biodiversity effects of seafood quantitatively comparable with the widely recognized relationship between biodiversity and productivity (26, 47, 52), and demonstrate a benefit of biodiversity for human nutritional well-being over and above the benefits of consuming a particular amount (biomass) or identity of aquatic species.

### Increasing seafood diversity increased contaminant exposure

We considered a range of trace elements, which, at high concentrations are known to be harmful to human health (57). We focused on four heavy metals considered as contaminants (methylmercury, cadmium, arsenic and lead) for which there exist public health guidelines for upper tolerable limits and for which seafood is potentially a major source of dietary intake (SI Appendix**, Figure S1, Table S2**). We examined the concentrations of these elements in the muscle tissues of 353 seafood species (thereby excluding parts such as viscera, liver and bones). We found that the same mechanisms that lead to a positive relationship between seafood species richness and nutritional benefits also contributed to exposure to a wider range of contaminants with higher species richness. We observed high levels of variation in contaminant concentrations across species (**Figure 4A-D**), and as we observed for micronutrients, these distributions were often right-skewed such that most species contained low contaminant concentrations and few contained high concentrations. On average, muscle tissue concentrations of contaminants were weakly positively correlated across species, such that species that contained high concentrations of one contaminant also contained high concentrations of another (average pairwise correlation = 0.09) (SI Appendix, **Figure S3**). When considering multiple contaminants simultaneously, increasing seafood species richness increased the number of contaminants which exceeded their upper tolerable limits (PTDI) in a 100g portion, referred to as *NC* (SI Appendix, Table S1 Equation A4), (*b_NC_* = 0.10, 95% CI 0.084, 0.12) (**Figure 4E**). When we considered the effects of biodiversity on exposure to each contaminant separately, we found that increasing species richness generally increased contaminant content per 100g, but the strength of this effect varied among contaminants. For example, increasing species richness from one to ten species was associated with doubling methylmercury concentrations on average, thereby reducing the maximum portion size before exceeding PTDI, referred to as *P_max_*, (SI Appendix, Table S1 Equation A3) (*b_Pmax_* = −0.25 95% CI −0.27, −0.22). For lead, however, the biodiversity effect was weaker (an order of magnitude smaller than for methylmercury). Increasing species richness from one to ten species was associated with only a 10% increase in average lead concentrations, and a small reduction in maximum portion size, *P_max_* (*b_Pmax_*= −0.039, 95% CI −0.049, −0.034). For cadmium, *b_Pmax_* = −0.12 (95% CI −0.13, −0.12), and for arsenic, *b_Pmax_* = −0.17 (95% CI −0.19, - 0.14). These differences in the strength of biodiversity effects among contaminants are linked to the shape of the distribution of their concentrations among fish species (i.e. normal vs. lognormal) (**Figure 4A-D**), and the more skewed the distribution, the stronger the biodiversity effect.

**Figure 4.**
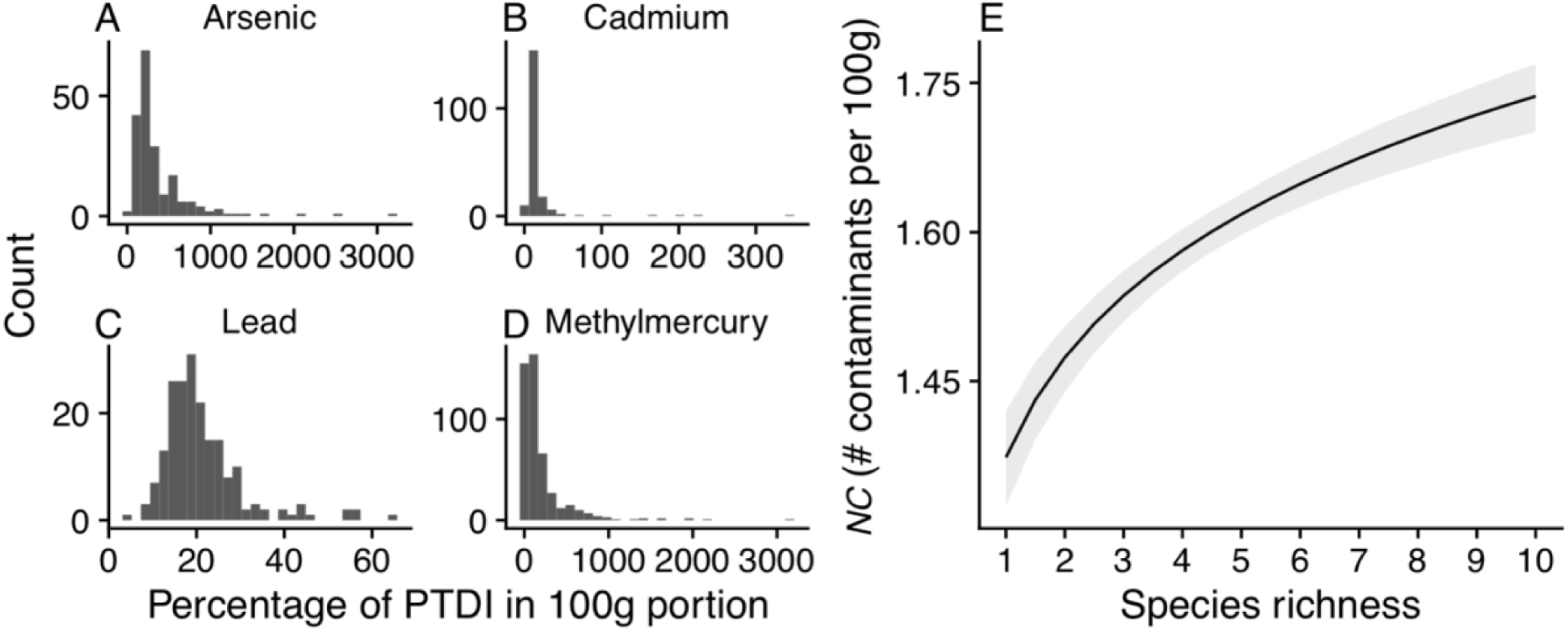
Frequency histograms of concentration of arsenic (**A**), cadmium (**B**), lead (**C**) and methylmercury (**D**) in edible muscle tissues of North American aquatic species relative to Provisional Tolerable Daily Intakes (PTDI). (**E)** Increasing seafood species richness increases the number of contaminants which exceed the upper tolerable limit (PTDI) in a 100g portion (*NC*). Black line indicates the mean *NC* from 1000 bootstrapped samples of seafood diets sampled from the North American seafood contaminant dataset and grey shading refers to 95% confidence intervals. Slope, *b_NC_* = 0.10, 95% CI 0.084, 0.12.

### Biodiversity benefits were consistent at local and global scales

Consistent with the positive biodiversity effects we observed when sampling diets from a global seafood species pool, we also found benefits of seafood diversity in a local context. We analyzed the effects of biodiversity on nutrient content in fourteen traditional indigenous North American diets of seafood harvested and consumed locally. We found a consistent, beneficial effect of biodiversity on *NT* and *P_min_*, although the magnitude of the biodiversity effect was generally lower at the local scale than the global scale (SI Appendix, **Figure S4**) (global *b_Pmin_* = −0.42 (95% CI −0.47, −0.38) vs mean local *b_Pmin_* = −0.25 ± 0.0091 S.E. and global *b_NT_* = 0.20 (95% CI 0.20, 0.21) vs mean local *b_NT_* = 0.097 ± 0.0082 S.E.). This finding is consistent with lower nutritional functional diversity (NFD, Table S1) *sensu* (34) (mean local *NFD* = 2.77 ± 0.17 S.E. vs. global *NFD* = 3.87) and higher nutritional functional evenness in local diets (mean local *NFEve* = 0.82 ± 0.0037 S.E. vs. global *NFEve* = 0.57) (SI Appendix, Methods section 3, SI Appendix, **Figure S5**), suggesting that functional consequences of changes to diversity in local seafood diets may be buffered by higher nutritional redundancy among species.

### Nutritional traits covary with ecological traits

We found that nutrient concentrations varied substantially among species in ways that differed for different nutrients (**Figure 2**) -- a condition necessary for biodiversity *per se* to increase nutritional benefits. The diversity that we observed in the nutrient content of edible portions (**Figure 2**) was partly explained by ecological attributes and functional traits: habitat, trophic position, body size, diet source and feeding mode (SI Appendix, Tables S4-S8). When considering all five micronutrients (calcium, iron, zinc, EPA and DHA) together, finfish, crustaceans and molluscs differed significantly in their multi-nutrient profiles (SI Appendix, Table S1, PERMANOVA, *F* _2,103_ = 3.429, *p* = 0.006). Among finfish, nutrient concentrations depended on which tissues were included in the edible portions (significant ‘body part’ effect shown in SI Appendix, Tables S4, S6-S11). Finfish species whose edible portions included organs such as liver or bones, had higher nutrient concentrations in the edible portion than those whose edible portions were restricted to muscle tissue (ANOVA *p* < 0.01 for calcium, iron and zinc concentrations; SI Appendix, **Figure S6**). Principal components analysis of multiple nutrient concentrations in species showed that essential element (calcium, iron and zinc) concentrations were typically negatively correlated with essential fatty acid concentrations (EPA and DHA) (SI Appendix, **Figure S7**), allowing complementarity among species to increase nutritional benefits. Specifically, high EPA and DHA concentrations traded off against low calcium and zinc concentrations (and vice-versa) (negative pairwise Pearson correlation coefficients; SI Appendix, **Figure S7**). When considering muscle tissues or muscle and skin tissues of finfish only (thereby eliminating the influence of body parts such as bones on nutrient concentration), concentrations of calcium, iron, zinc, EPA and DHA were associated with ecological traits across species, including habitat and diet source (e.g. demersal vs. pelagic), body size and trophic position (**Figure 5**). Relationships between species’ nutrient tissue concentrations and their habitats and trophic positions have been predicted by ecological stoichiometry theory (58). Relationships between ecological traits and nutrient concentrations differed for different nutrients. For example, tissue concentrations decreased with body size for calcium, but not for the other microelements, EPA or DHA (**Figure 5**, SI Appendix, Figures S8-S10, Tables S4 - S8). Species at lower trophic positions had higher zinc concentrations in their tissues than species at higher trophic positions, but we did not observe this relationship for other nutrients (SI Appendix, Tables S4 - S8). These examples illustrate that trade-offs and variation in nutrient concentrations across species were associated with variation in different ecological traits and roles that species play in ecosystems.

**Figure 5.**
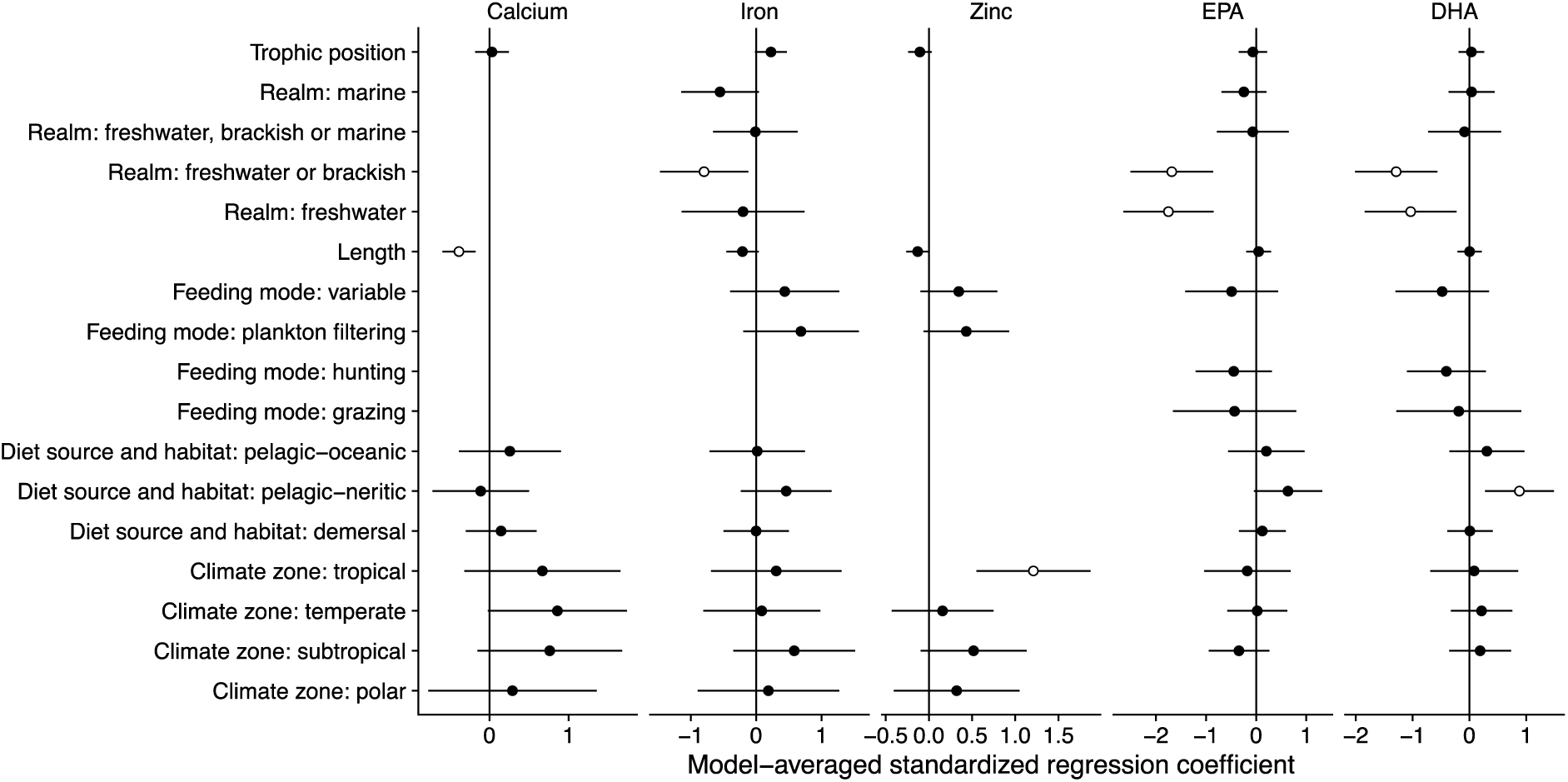
Nutrient concentrations in finfish muscle tissue vary with ecological traits in ways that differ among the essential trace elements (calcium, iron, zinc) and the essential fatty acids (EPA and DHA). Model-averaged standardized regression coefficients and 95% confidence intervals from phylogenetic least squares regression (see SI Appendix, Methods 4 for full model description) are shown for samples including muscle or muscle and skin tissues only. Traits for which there is no symbol did not appear in any of the models in the top 95% set (with cumulative sum of Akaike weights <= 0.95). Open symbols indicate coefficient estimates for which 95% confidence intervals do not encompass zero. Note that x-axes differ across panels for clarity of presentation. Number of species: *n* = 155 for EPA, *n* = 159 for DHA, *n* = 104 for calcium, *n* = 99 for iron, and *n* = 90 for zinc. For model results including tissues other than muscle tissue, see SI Appendix, Tables S4-S11.

### Nutritional value was linked to diversity of ecological functions

Ecological functional diversity of a species assemblage captures variation in traits and ecological roles of species (48, 49), and is understood to play an important role in the relationship between biodiversity and ecosystem function (**Figure 6A**). We assessed the relationship between ecological functional diversity of seafood species diets (independent from species richness) and nutritional value. Consistent with observations for ecosystem functions such as productivity and biomass, nutritional benefits and ecological functional diversity were positively related, such that seafood diets with higher ecological functional diversity also provided higher nutritional value (i.e. were more likely to reach five micronutrient RDA targets simultaneously, *NT* = 5) (**Figure 6B**). Ecological functional diversity increased with species richness (SI Appendix, **Figure S11**), and higher levels of ecological functional diversity were also associated with lower minimum portion size required (SI Appendix, **Figure S12**). Because aquatic assemblages with higher ecological functional diversity have been shown to exploit more diverse resources, transform and transport energy and materials more efficiently, produce higher yields, and be more productive and resilient over time (42, 59–61), it is possible that the provisioning of multiple micronutrients occurs in tandem with a range of other ecological functions.

**Figure 6.**
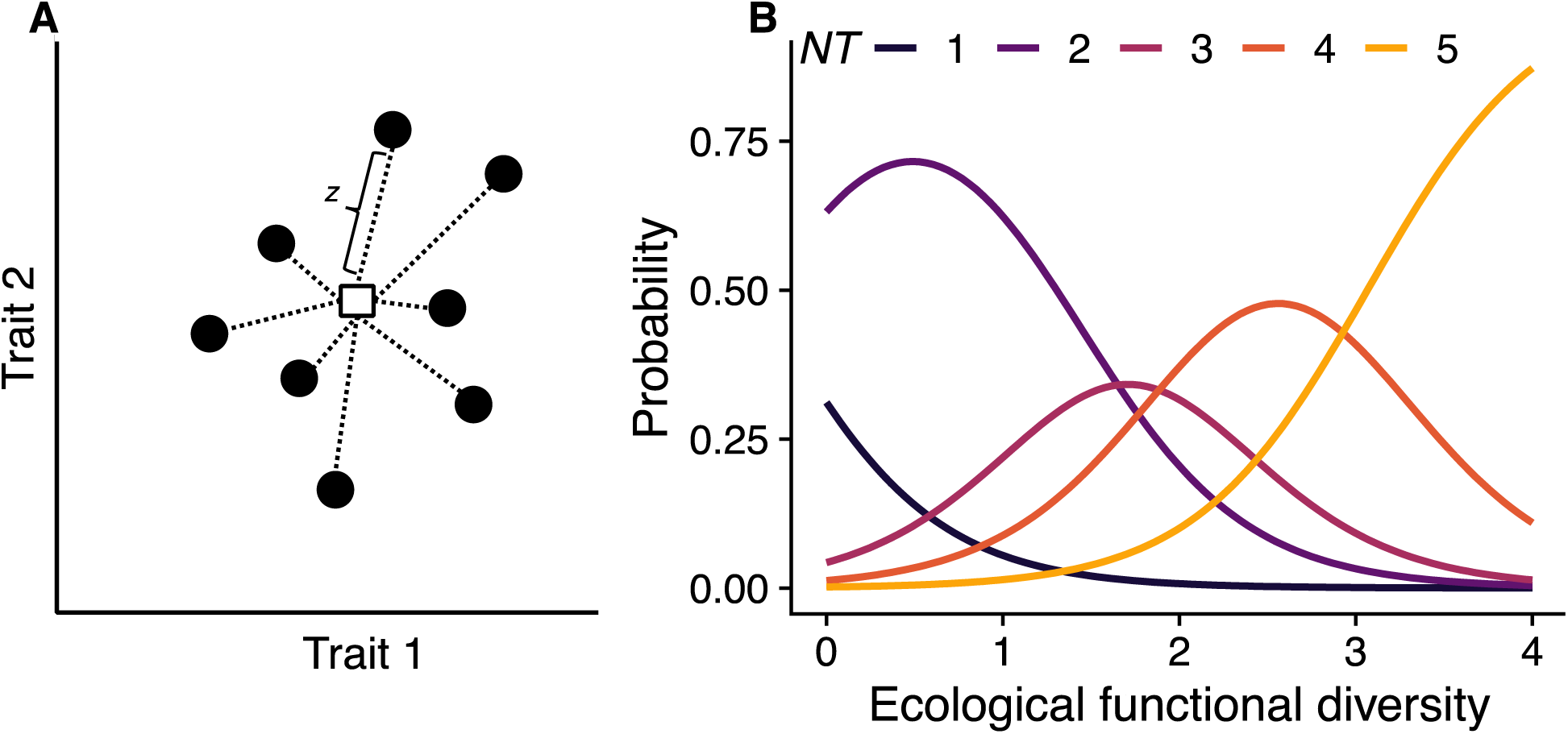
**A**) Ecological functional diversity captures variation in the ecological traits and ecological roles of species, and can be quantified as the average of the distances between species’ trait values (circles) and the centre of functional trait space (square) (indicated by *z*) (adapted from (93)). **B**) Seafood diets with higher levels of ecological functional diversity are associated with higher levels of nutritional benefits (i.e. higher number of RDA targets per 100g portion). Probability of reaching five micronutrient RDA targets simultaneously in a single 100g portion (*NT* = 5, light orange line) increases as the ecological functional diversity of the diet increases. Probabilities predicted from ordinal logistic regression (log odds of functional diversity = 2.08, 95% CI (0.52, 3.64), *n* = 1000 resampled diets).

## Discussion

Biodiversity of seafood provides high levels of nutritional benefits to humans because nutrient concentrations vary substantially among species in ways that differ for different nutrients (**Figures 2, 5**, SI Appendix, **Figure S7**). The effects of species richness observed for nutritional benefits equal or exceed mean observed diversity effects documented for plant and forest species richness and productivity (12, 40, 52, 62). Our findings build on evidence showing that biodiversity in marine systems enhances ecological functions by extending this paradigm to human nutrition. In agro-ecosystems, specific combinations of species such as corn, pumpkin and beans have been planted to exploit complementary traits (shade provisioning, nitrogen fixation and biomass production), to attain higher yields, more resilient crops and enhanced nutritional benefits (35). By demonstrating that nutritional benefits of seafood can be understood as a consequence of seafood species richness and ecological functional diversity, we have shown that diverse seafood assemblages also provide nutritional and other ecological benefits simultaneously.

Our analysis provides new and robust evidence that biodiversity is critical to multifunctionality of ecosystem services when function thresholds are grounded *a priori* in multivariate metrics meaningful for human well-being such as RDA. Our approach overcomes the critique that multifunctionality is not enhanced by biodiversity (63), but rather a statistical artifact of how multifunctionality was commonly estimated. Our findings are robust to a range of RDA target levels (SI Appendix, **Figure S2**). In the case of *P_min_*, we found that the benefit of biodiversity was consistent across all RDA target levels considered, and in the case of *NT*, we found beneficial effects of biodiversity across a range levels of nutritional value that are significant for human nutrition (i.e. 1-40% RDA per 100g portion), highlighting the importance of species richness in seafood diets. More generally, ecosystem service benefits, as defined in metrics of human wellbeing rather than the traits of the species pool under consideration (e.g., biomass or stability of the food web), typically are produced by several underlying ecosystem functions (54). The strong effects of diversity on multifunctional benefits observed here may also apply to relationships between diversity and other services e.g., desired filtration rates of pollutants in wetlands (64), or desired pest consumption rates in agricultural systems (65).

### Physiological and ecological mechanisms underlying nutritional benefits and risks

When comparing the magnitude of the biodiversity-ecosystem function scaling parameter, *b*, among the nutrients and contaminants, we found that the magnitude of *b* was higher for nutrients with more skewed distributions of tissue concentrations (**Figure 2, Figure 4**). For all of these highly skewed distributions, we observed a strong biodiversity effect (i.e. high values of *b*). However, for protein and lead in muscle tissues, we observed symmetrical tissue concentration distributions. In these two cases, the biodiversity effect was either non-existent (in the case of protein) or very weak (in the case of lead). This finding suggests that biodiversity effects may increase when the trait distribution is skewed and includes some species with extreme trait values, known as ‘functionally specialized’ and ‘functionally unique’ species (9, 66, 67). Understanding the drivers of the tissue concentration distribution among species may lend insights into how biodiversity effects may change as the nutrient under consideration changes or as the environment changes.

Increasing seafood species richness increases nutritional benefits as well as contaminant exposure. Increasing biodiversity can have negative consequences for some ecosystem functions, despite positive consequences for others. On balance, the benefits of diversity may outweigh the negative effects on function when considering multiple functions together (68). Here we found that the same mechanisms (lognormal and complementary tissue concentration distributions among species) that contributed to the positive relationship between biodiversity and human nutritional benefits also applied to contaminant exposure. As a result, increasing seafood species richness comes with both benefits and risks (69–71). However, interpreting our results in the context of public health outcomes is complicated by the fact that epidemiological evidence on health outcomes of contaminant exposure is mixed and likely dependent on complex social and health risks. The health risks of contaminant exposure depend on other health factors such as smoking or nutrients in the diet (72, 73) and disease status. Complex interactions among multiple diet and health risk factors (74) were beyond the scope of this study, but would be necessary to understand exposure risks from seafood consumption in any particular community. Nonetheless, our results suggest that while increasing biodiversity increases contaminant exposure, it also increases nutritional benefit, and reduces the portion sizes required to meet nutritional demands. Finding a balance between seafood biodiversity, seafood biomass consumption, and the resulting risks and benefits will be critical for both human and ecosystem health.

### Aquatic biodiversity and food security in a changing world

Seafood-derived nutrition plays an important role in food security. The link that we have demonstrated between seafood biodiversity (species richness and ecological functional diversity) and nutrition in an ecological framework unites three of the United Nations Sustainable Development Goals focused on biodiversity, hunger and well-being (75). More than two billion people suffer from micronutrient deficiencies (76, 77), and many of the most nutritionally vulnerable populations – those that are deficient in essential micronutrients during particularly sensitive stages of life (i.e. pregnancy, breastfeeding and childhood) - may rely heavily on local aquatic ecosystems to meet their nutritional demands (15, 19, 22, 78). These populations may have access to a limited amount of locally available seafood tissue each day, suggesting that nutritional efficiency (i.e. lower *P_min_*) provided by biodiversity in wild-caught seafood may be particularly important for these populations. Regions of high nutritional vulnerability continue to experience major changes in biodiversity and ecosystem structure (6, 17, 79), and climate change is compounding threats to the sustainability of capture fisheries (2, 80). Our results suggest that declines in these aspects of diversity in wild aquatic ecosystems could make achieving sustainability goals for food security via seafood even more difficult. Seafood diets composed of more species, and groups of species with higher levels of ecological functional diversity, are more likely to provide more nutrients per unit biomass than less diverse seafood diets, while also maintaining high levels of ecosystem function (26). This finding bridges the growing understanding of hidden hunger and food security with the large theoretical and empirical understanding of relationships between biodiversity, ecosystem function and benefits to people. Biodiversity in natural aquatic systems can be maintained by reducing pollution and overharvest and by allowing ecosystems to adapt to climate change, and these measures could also benefit humanity directly through seafood provisioning.

## Conclusions

Nutritional value appears to be derived from ecological diversity, suggesting links between the complexity of aquatic ecosystems and their capacity to produce nutritional benefits. While the role of seafood is well-recognized as an important source of protein in the human diet, the role of seafood biodiversity as an important aspect of the provision of essential micronutrients has been overlooked. Our results reveal that aspects of ecological structure including species and ecological functional diversity enhance nutritional benefits, while also increasing contaminant exposure, thereby linking the processes that structure ecosystems with their potential benefits and risks to human nutrition and health.

## Methods

To test our hypotheses about aquatic species diversity and potential nutritional benefits for human well-being, we assembled a new database of nutritional values by synthesizing observations from existing data (SI Appendix Methods, **Figure S1**). To build the database, we identified quantitative and comparable measures of nutritional content for the edible portions of aquatic species (SI Appendix Methods 1.1). We then identified metrics for relating nutrient content to human health (Methods 1.1, 1.3). We repeated this exercise for contaminants (Methods 1.2). Next, we adapted an approach from biodiversity - ecosystem functioning theory for quantitatively assessing the potential nutritional value of seafood diets varying in their species composition and diversity for multiple nutrients separately and simultaneously (Methods 2.1). To test predictions of biodiversity-ecosystem functioning theory, we simulated potential diets at global and local scales and fit models to test for effects of species richness and functional diversity (Methods 2.2 - 2.3). SI Appendix, **Figure S1** provides a graphical overview of our analyses.

### 1. Metrics

#### 1.1 Quantifying nutritional value for single nutrients

We characterized an aquatic species’ nutritional value by drawing on two well-established nutritional metrics: nutrient concentration (mg/100 g edible portion) and Recommended Dietary Allowances (RDA). RDAs are developed following health guidelines to quantify the recommended amount of a particular nutrient required to maintain health (50). We used RDAs established by the Food and Nutrition Board of the United States Institute of Medicine to quantify the daily intake level of a nutrient needed to meet the requirements of 97–98% of healthy adults (females aged 19-50) (50) (SI Appendix, Table S2). We refer to an RDA-defined threshold as an *RDA target* (SI Appendix, Tables S1 and S2). We calculated a ratio of the nutrient content in a 100g edible portion relative to the RDA (or fraction thereof) for that nutrient. For many species, nutrient concentrations in edible tissues provided only small fractions of the RDA (SI Appendix, Table S2). Following the IOM, we chose an RDA target as 10% of RDA for a given nutrient in a single portion (SI Appendix, Table S2) because this is a minimum threshold for a food to be considered of nutritional benefit (50). We defined the minimum portion size required, *P_min_*, as the minimum amount of edible seafood issue (g) required to reach the RDA target for a given nutrient (or a set of nutrients, Methods 1.3) (SI Appendix, Table S1, Equation A1). We quantified the sensitivity of our biodiversity findings to different threshold levels of RDA for both single nutrient analyses and multi-nutrient analyses (Methods 2.2) and found that for *P_min_*, they did not vary with threshold level, and for *NT*, they varied with threshold, but were robust to RDA threshold levels between 1% and 40% RDA (SI Appendix, **Figure S2**).

We examined data on concentrations of macronutrients including protein and fat, as well as five micronutrients (*n* = 5041 observations of nutrient concentrations, SI Appendix, **Figure S1A**): metals beneficial at low concentrations but toxic at high concentrations (zinc and iron), one beneficial mineral (calcium) and the polyunsaturated fatty acids eicosapentaenoic acid (EPA) and docosahexaenoic acid (DHA). We chose these five micronutrients because we required that RDA standards exist for each one (50) and that each nutrient has known functions in organismal physiology, and are considered ‘biologically essential’ because they are required by organisms to grow or reproduce. The concentration of biologically essential nutrients in organisms’ tissues are controlled homeostatically by organisms (81, 82), and therefore might be biologically related to ecological trait values we considered in our trait analysis.

#### 1.2 Quantifying contaminant exposure in terms of human health risks

We characterized an aquatic species’ contaminant content relative to established public health guidelines for exposure. We used the Provisional Tolerable Weekly Intake (PTWI) developed by the FAO/WHO Expert Committee on Food Additives (JEFCA) (57), which estimates the amount of a substance in air, food, soil or drinking water that can be assimilated weekly per unit body weight over a lifetime without appreciable health risk (57). To make this metric comparable to RDA targets defined above, we quantified contaminant exposure per day by dividing the PTWI by seven and using the body weight of a 70-kg person, to calculate a daily tolerable limit for use in our analyses, which we refer to as Provisional Tolerable Daily Intake, PTDI (SI Appendix, Table S1). In our analyses, we consider a tolerable upper limit to be exceeded if an edible portion contains 100% or more of the PTDI. We examined the sensitivity of this choice of 100% threshold by considering lower values (e.g. 50%) and found that for all values between 50% and 100% of PTDI our findings remain consistent.

#### 1.3 Defining nutritional benefits and risks for multiple nutrients or contaminants across diverse species groups

To test our hypothesis that nutritional value may depend on seafood diversity in a way that considers multiple nutrients at the same time, we used two metrics that considered multiple micronutrients simultaneously (**Figure 1**): (i) the minimum portion size introduced above, *P_min_*, (SI Appendix, Table S1, Equation A1), which quantifies the amount of tissue, in grams, required to reach the RDA target for 5 nutrients simultaneously and (ii) number of nutrients for which RDA targets are met in a standard 100g edible portion, *NT* (SI Appendix, Table S1, Equation A2). Note that our *NT* differs from another measure ‘Nutritional functional diversity, NFD’ (34) which is a measure of nutritional profile in multivariate trait space and does not explicitly quantify the number of nutrient RDA targets met in a portion. Our *NT* is a measure of potential nutritional value and is derived from the biodiversity-ecosystem function perspective to allow us to compare nutritional value with other functions that depend on biodiversity (e.g., productivity, resource cycling). This measure does not consider the potential physiological interactions or co-benefits of consuming multiple nutrients that would be considered in metabolic models for human health. To test our hypothesis that contaminant exposure also varies with seafood diversity, we defined *P_max_*, as the amount of tissue beyond which a tolerable upper limit (PTDI) for a given contaminant would be reached (SI Appendix, Table S1, Equation A3, SI Appendix Methods 2). Analogous to the *NT* metric defined above for the nutrients, we defined the number of distinct PTDIs in a 100g portion as *NC* (Table S1, Equation A4, SI Appendix Methods 2). We chose to standardize *NT* and *NC* relative to 100g of seafood per day because 100g edible portion is a standard metric used in the food and nutrition literature (37, 83) and consumption rates of 100g per day is within the range of daily consumption rates for many communities that rely heavily on seafood to meet nutritional needs and for whom seafood may also pose substantial contaminant exposure risks (16, 45, 84, 85).

### 2. Statistical analyses and hypothesis testing

#### 2.1 Modeling the biodiversity effect for sample diets

To quantitatively compare effects of biodiversity among possible seafood diets varying in species richness, we modeled species richness effects as a power function:

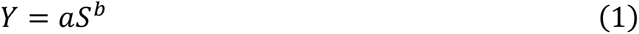

where the parameter *b* is referred to as the ‘biodiversity effect’ and describes the relationship between a change in species richness, *S*, and a measure of function, *Y*, such as *P_min_* or *NT*, where *a* is a constant (12, 52).

#### 2.2 Simulating seafood diets and estimating nutritional benefits at different seafood species richness levels

We tested the effect of species richness, *S*, on nutritional value by randomly assembling diets from the global seafood species pool at varying levels of species richness (SI Appendix, **Figure S1B**). In our analyses of *NT*, we kept the total biomass constant at each level of species richness (100g). In analyses using the global dataset (547 species) (SI Appendix Methods 1.1), we assembled diets from the entire global species pool, choosing species at random without replacement. This way of simulating diets certainly ignores economic, social and cultural factors that affect which species people consume, but allows us to consider the potential effect of biodiversity on diets before diets are filtered by these other processes. To assess potential effects of biodiversity on nutritional value for populations that consume seafood locally from a restricted species pool, we sampled diets from species contained within traditional diets in fourteen indigenous cultures in North America (SI Appendix, Methods 1.4).

We created sample diets by sampling 10 species at random from the global species pool and then assembling seafood diets from all possible combinations of these ten randomly chosen species at 10 levels of species richness (1–10) to generate 1023 simulated diets (SI Appendix, **Figure S1B, i**). At each level of species richness, we assembled diets following the typical experimental design employed to test the hypothesis that biodiversity affects ecosystem functioning, analogous to a biodiversity-ecosystem function experiment with a replacement design (86), where species’ abundances in the diet decline proportionally as species richness increases such that each species contributed an equal proportion of biomass. For each diet at each level of species richness (i.e. *n* = 1023), we calculated *P_min_*, (Methods 1.1, 1.3, SI Appendix, Table S1 Equation A1, SI Appendix, **Figure S1C, ii**) (for either: one of six possible nutrient targets individually (protein, calcium, iron, zinc, EPA and DHA), or five micronutrients (calcium, iron, zinc, EPA and DHA targets simultaneously) (Methods 1.3). To estimate *NT* (Methods 1.3, SI Appendix, **Figure S1C, ii**), we quantified the number of distinct nutrient RDA targets by assigning each diet (*n* = 1023) a set of 0’s or 1’s according to whether that combination met the RDA target for each nutrient (see SI Appendix, Table S1, Equation A2). This approach allowed us to explore how likely it would be for potential human diets containing different numbers of seafood species to reach RDA targets for a given number of micronutrients (*NT* ranges between 0 and 5), assuming that seafood species were included in the human diet at different levels of richness at random. At each level of species richness, we averaged *P_min_* and *NT*. We then repeated this process of random sampling ten species from the species pool, assembling diets at each level of richness, estimating metrics of nutritional value and averaging at each richness level 1000 times, yielding 1000 estimates of each metric at each richness level (SI Appendix, **Figure S1C, iii**).

#### 2.3 Testing hypotheses that biodiversity enhances nutritional benefits

We tested the hypothesis that complementarity in nutrient concentrations among species increases nutritional benefits by increasing *NT* (SI Appendix, **Figure S1D**). We quantified the effect of seafood species richness, *S*, on *NT* at each richness level estimated in Methods 2.2 (*n* = 1000 estimates of *NT* per richness level), by fitting a power function of the form shown in Equation 1,

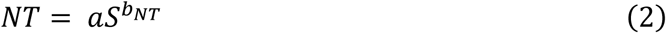

where the parameter *b_NT_* describes the relationship between a change in species richness, *S*, and a change in *NT*, and *a* is a constant.

We tested the hypothesis that complementarity in nutrient concentrations among species reduces the minimum portion size required, *P_min_* (Methods 1.1, 1.3, 2.2), by estimating the effect of species richness, *S*, on *P_min_* (*n* = 1000 estimates of *P_min_* per richness level) at each richness level using

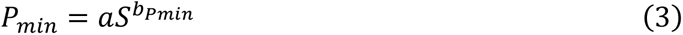

where the parameter *b_Pmin_* describes the relationship between a change in species richness, *S*, and a change in *P_min_*, and *a* is a constant (SI Appendix, **Figure S1D**).

We estimated *a* and *b* in Equations 2 and 3 using non-linear regression using the *nls.LM* function in the *minpack.lm* package in R (87). We conducted all analyses in R version 3.3.2 (88). To quantify uncertainty in parameter estimates associated with the fit of Equations 2 and 3 to our estimates of function (Methods 2.2), we calculated bootstrapped confidence (*n* = 1000 bootstraps) intervals using non-parametric bootstrapping of mean centered-residuals using the *nlsBoot* function in the R package *nlstools* (89). For both *P_min_* and *NT*, we tested the hypothesis that biodiversity enhances nutritional benefits by assessing whether the estimate of the scaling exponent, *b*, had confidence intervals not overlapping zero.

We tested the hypothesis that nutrient concentrations are related to species’ ecological traits in two ways: 1) testing whether multi-nutrient profiles (i.e. concentrations of all five micronutrients) differ among major phylogenetic groups using PERMANOVA (SI Appendix, Methods 4) and 2) whether differences in single nutrient concentrations differ with species’ ecological traits using phylogenetic least squares regression (SI Appendix, Methods 4). We quantified the relationship between nutritional benefits and ecological functional diversity (EFD, SI Appendix, Table S1) by estimating *NT*, *P_min_* and *EFD*, estimated as functional dispersion, of diets simulated from the global species pool (SI Appendix, Methods 5).

#### 2.4. Uncertainties

There are several sources of uncertainty in our analyses. First, there are substantial sources of uncertainty in food composition estimates. The data in our dataset meet international standards for data quality and standardization, meaning that we followed guidelines for checking food composition data and converting units, denominators and expressions (90). Still, tissue concentrations may vary depending on analytical techniques, labs, season, diet of the animal, life stage etc. Some of these sources of uncertainty (e.g. differences in analytical techniques) are unavoidable consequences of synthesizing previously published data collected across many labs. We assumed that these uncertainties in the data were randomly distributed over our geographically and taxonomically diverse dataset. Further uncertainty is associated with how well our set of 547 species represents the global pool of seafood consumed. We do not know whether our sample is random or biased, though we can say that our dataset includes 41 of the 67 most consumed species worldwide (as determined by FAO production volumes (91), species with capture production of 150 000 tonnes or more, after removing species for which the majority of production volume is diverted to fish meal and oil (92), SI Appendix, Table S13). A remaining source of variation among samples is likely due to natural sources of variation associated with seasonal and other sources of temporal variability, which we consider to be an important component of biodiversity.

## Acknowledgments

We thank P. Thompson, A. Gehman, W. Cheung, M. Ruckelshaus and C. Harley, M. Deith, L. Dee, and F. Isbell for helpful comments on previous versions of this manuscript. Portions of this manuscript were adapted from and based on work contained in the doctoral thesis of J.R. Bernhardt from the University of British Columbia.

## Data and materials availability

Data and code are available at https://github.com/JoeyBernhardt/Nutrient_analysis, archived at 10.5281/zenodo.4474988. Data are available on Dryad (https://doi.org/10.5061/dryad.rn8pk0p8t).

## References

1. R. Elahi, et al., Recent trends in local-scale marine biodiversity reflect community structure and human impacts. Curr. Biol. 25, 1938–1943 (2015).

2. M. L. Pinsky, B. Worm, M. J. Fogarty, J. L. Sarmiento, S. A. Levin, Marine taxa track local climate velocities. Science 341, 1239–1242 (2013).

3. D. J. McCauley, et al., Marine defaunation: animal loss in the global ocean. Science 347, 1255641 (2015).

4. FAO, The State of the World’s Biodiversity for Food and Agriculture, J. Bélanger, D. Pilling, Eds. (FAO Commission on Genetic Resources for Food and Agriculture Assessments, 2019).

5. IPBES, Global assessment report on biodiversity and ecosystem services of the Intergovernmental Science-Policy Platform on Biodiversity and Ecosystem Services, E. S. Brondizio, J. Settele, S. Díaz, H. T. Ngo, Eds. (2019).

6. G. P. Jones, M. I. McCormick, M. Srinivasan, J. V. Eagle, Coral decline threatens fish biodiversity in marine reserves. Proc. Natl. Acad. Sci. U. S. A. 101, 8251–8253 (2004).

7. R. A. Myers, B. Worm, Rapid worldwide depletion of predatory fish communities. Nature 423, 280–283 (2003).

8. J. A. Hutchings, R. A. Myers, What Can Be Learned from the Collapse of a Renewable Resource? Atlantic Cod, Gadus morhua, of Newfoundland and Labrador. Can. J. Fish. Aquat. Sci. 51, 2126–2146 (1994).

9. C. Pimiento, et al., Functional diversity of marine megafauna in the Anthropocene. Sci Adv 6, eaay7650 (2020).

10. S. S. Myers, et al., Human health impacts of ecosystem alteration. Proc. Natl. Acad. Sci. U. S. A. 110, 18753–18760 (2013).

11. F. Isbell, et al., Linking the influence and dependence of people on biodiversity across scales. Nature 546, 65–72 (2017).

12. M. I. O’Connor, et al., A general biodiversity--function relationship is mediated by trophic level. Oikos 126, 18–31 (2017).

13. E. Chivian, “Biodiversity: Its Importance to Human Health” (Center for Health and the Global Environment Harvard Medical School, 2002).

14. FAO, The State of Food Security and Nutrition in the World: Building Resilience for Peace and Food Security, Rome: Food and Agriculture Organization of the United Nations, Ed. (2017).

15. H. V. Kuhnlein, O. Receveur, Local Cultural Animal Food Contributes High Levels of Nutrients for Arctic Canadian Indigenous Adults and Children. J. Nutr. 137, 1110–1114 (2007).

16. A. M. Cisneros-Montemayor, D. Pauly, L. V. Weatherdon, Y. Ota, A Global Estimate of Seafood Consumption by Coastal Indigenous Peoples. PLoS One 11, e0166681 (2016).

17. J. G. Molinos, et al., Climate velocity and the future global redistribution of marine biodiversity. Nat. Clim. Chang. 6, 83–88 (2016).

18. WHO, Connecting global priorities: biodiversity and human health: a state of knowledge review, World Health Organization and Secretariat of the Convention on Biological Diversity, Ed. (2015).

19. C. D. Golden, et al., Fall in fish catch threatens human health. Nature 534, 317–320 (2016).

20. B. Belton, S. H. Thilsted, Fisheries in transition: Food and nutrition security implications for the global South. Global Food Security 3, 59–66 (2014).

21. FAO, The State of World Fisheries and Aquaculture 2018 - Meeting the sustainable development goals (2018).

22. N. Kawarazuka, C. Béné, Linking small-scale fisheries and aquaculture to household nutritional security: an overview. Food Security 2, 343–357 (2010).

23. B. J. Cardinale, et al., Biodiversity loss and its impact on humanity. Nature 486, 59–67 (2012).

24. D. Tilman, et al., The Effects of Plant Composition and Diversity on Ecosystem Processes. Science 277, 1302–1305 (1997).

25. M. Loreau, Biodiversity and ecosystem functioning: recent theoretical advances. Oikos 91, 3–17 (2000).

26. J. E. Duffy, C. M. Godwin, B. J. Cardinale, Biodiversity effects in the wild are common and as strong as key drivers of productivity. Nature 549, 261–264 (2017).

27. M. T. Ruel, Operationalizing dietary diversity: a review of measurement issues and research priorities. J. Nutr. 133, 3911S–3926S (2003).

28. E. A. Frison, J. Cherfas, T. Hodgkin, Agricultural Biodiversity Is Essential for a Sustainable Improvement in Food and Nutrition Security. Sustain. Sci. Pract. Policy 3, 238–253 (2011).

29. J. Fanzo, D. Hunter, T. Borelli, F. Mattei, Diversifying Food and Diets: Using Agricultural Biodiversity to Improve Nutrition and Health (Routledge, 2013).

30. E. A. Frison, I. F. Smith, T. Johns, J. Cherfas, P. B. Eyzaguirre, Agricultural biodiversity, nutrition, and health: making a difference to hunger and nutrition in the developing world. Food Nutr. Bull. 27, 167–179 (2006).

31. Á. Toledo, B. Burlingame, Biodiversity and nutrition: A common path toward global food security and sustainable development. J. Food Compost. Anal. 19, 477–483 (2006).

32. A. D. Jones, Critical review of the emerging research evidence on agricultural biodiversity, diet diversity, and nutritional status in low- and middle-income countries. Nutr. Rev. 75, 769–782 (2017).

33. C. Lachat, et al., Dietary species richness as a measure of food biodiversity and nutritional quality of diets. Proc. Natl. Acad. Sci. U. S. A. 115, 127–132 (2018).

34. R. Remans, et al., Assessing nutritional diversity of cropping systems in African villages. PLoS One 6 (2011).

35. F. A. J. DeClerck, J. Fanzo, C. Palm, R. Remans, Ecological approaches to human nutrition. Food Nutr. Bull. 32, S41–50 (2011).

36. D. Penafiel, C. Lachat, R. Espinel, P. Van Damme, P. Kolsteren, A systematic review on the contributions of edible plant and animal biodiversity to human diets. EcoHealth 8, 381–399 (2011).

37. R. Remans, S. A. Wood, N. Saha, T. L. Anderman, R. S. DeFries, Measuring nutritional diversity of national food supplies. Global Food Security 3, 174–182 (2014).

38. S. A. Wood, Nutritional functional trait diversity of crops in south-eastern Senegal. J. Appl. Ecol. 55, 81–91 (2018).

39. M. Loreau, A. Hector, Partitioning selection and complementarity in biodiversity experiments. Nature 412, 72–76 (2001).

40. P. B. Reich, et al., Impacts of biodiversity loss escalate through time as redundancy fades. Science 336, 589–592 (2012).

41. B. J. Cardinale, Biodiversity improves water quality through niche partitioning. Nature 472, 86–89 (2011).

42. L. E. Dee, et al., Functional diversity of catch mitigates negative effects of temperature variability on fisheries yields. Proc. Biol. Sci. 283 (2016).

43. R. Hilborn, T. P. Quinn, D. E. Schindler, D. E. Rogers, Biocomplexity and fisheries sustainability. Proc. Natl. Acad. Sci. U. S. A. 100, 6564–6568 (2003).

44. S. Wang, M. Loreau, Biodiversity and ecosystem stability across scales in metacommunities. Ecol. Lett. 19, 510–518 (2016).

45. J. R. Bogard, et al., Higher fish but lower micronutrient intakes: Temporal changes in fish consumption from capture fisheries and aquaculture in Bangladesh. PLoS One 12 (2017).

46. C. C. Hicks, et al., Harnessing global fisheries to tackle micronutrient deficiencies. Nature 574, 95–98 (2019).

47. D. Tilman, The Ecological Consequences of Changes in Biodiversity: A Search for General Principles. Ecology 80, 1455 (1999).

48. D. Tilman, Functional diversity. Encyclopedia of biodiversity 3, 109–120 (2001).

49. C. Violle, et al., Let the concept of trait be functional! Oikos 116, 882–892 (2007).

50. IOM, “Dietary reference intakes for calcium and vitamin d” (2011).

51. F. Micheli, B. S. Halpern, Low functional redundancy in coastal marine assemblages. Ecol. Lett. 8, 391–400 (2005).

52. B. J. Cardinale, et al., The functional role of producer diversity in ecosystems. Am. J. Bot. 98, 572–592 (2011).

53. J. E. K. Byrnes, et al., Investigating the relationship between biodiversity and ecosystem multifunctionality: Challenges and solutions. Methods in Ecology and Evolution 5, 111–124 (2014).

54. P. Manning, et al., Redefining ecosystem multifunctionality. Nature Ecology & Evolution 2, 427–436 (2018).

55. A. Hector, R. Bagchi, Biodiversity and ecosystem multifunctionality. Nature 448, 188–190 (2007).

56. L. Gamfeldt, H. Hillebrand, P. R. Jonsson, Multiple functions increase the importance of biodiversity for overall ecosystem functioning. Ecology 89, 1223–1231 (2008).

57. WHO, Evaluation of Certain Contaminants in Food: Seventy-third Report of the Joint FAO/WHO Expert Committee on Food Additives (2011).

58. R. W. Sterner, J. J. Elser, Ecological stoichiometry: the biology of elements from molecules to the biosphere (Princeton University Press, 2002).

59. J. S. Lefcheck, et al., Tropical fish diversity enhances coral reef functioning across multiple scales. Sci Adv 5, eaav6420 (2019).

60. C. Mora, et al., Global human footprint on the linkage between biodiversity and ecosystem functioning in reef fishes. PLoS Biol. 9, e1000606 (2011).

61. J. S. Lefcheck, J. E. Duffy, Multitrophic functional diversity predicts ecosystem functioning in experimental assemblages of estuarine consumers (2015) https:/doi.org/10.1890/14-1977.1.

62. J. Liang, et al., Positive biodiversity-productivity relationship predominant in global forests. Science 354, aaf8957 (2016).

63. L. Gamfeldt, F. Roger, Revisiting the biodiversity--ecosystem multifunctionality relationship. Nature Ecology & Evolution 1, s41559–017 (2017).

64. T. Boyer, S. Polasky, Valuing urban wetlands: A review of non-market valuation studies. Wetlands 24, 744–755 (2004).

65. D. S. Karp, et al., Forest bolsters bird abundance, pest control and coffee yield. Ecol. Lett. 16, 1339–1347 (2013).

66. D. Mouillot, N. A. J. Graham, S. Villéger, N. W. H. Mason, D. R. Bellwood, A functional approach reveals community responses to disturbances. Trends Ecol. Evol. 28, 167–177 (2013).

67. R. P. Leitao, et al., Rare species contribute disproportionately to the functional structure of species assemblages. Proceedings of the Royal Society B: Biological Sciences 283, 20160084 (2016).

68. J. S. Lefcheck, et al., Biodiversity enhances ecosystem multifunctionality across trophic levels and habitats. Nat. Commun. 6, 6936 (2015).

69. M. C. Sheehan, et al., Global methylmercury exposure from seafood consumption and risk of developmental neurotoxicity: a systematic review. Bull. World Health Organ. 92, 254–269F (2014).

70. H. V. Kuhnlein, H. M. Chan, Environment and contaminants in traditional food systems of northern indigenous peoples. Annu. Rev. Nutr. 20, 595–626 (2000).

71. P. W. Davidson, et al., Methylmercury and neurodevelopment: longitudinal analysis of the Seychelles child development cohort. Neurotoxicol. Teratol. 28, 529–535 (2006).

72. D. Mozaffarian, E. B. Rimm, Fish intake, contaminants, and human health: evaluating the risks and the benefits. JAMA 296, 1885–1899 (2006).

73. D. Kinloch, H. Kuhnlein, D. C. Muir, Inuit foods and diet: a preliminary assessment of benefits and risks. Sci. Total Environ. 122, 247–278 (1992).

74. M. O. Gribble, et al., Mercury, selenium and fish oils in marine food webs and implications for human health. J. Mar. Biol. Assoc. U. K. 96, 43–59 (2016).

75. UN, Transforming our world : the 2030 Agenda for Sustainable Development, UN General Assembly, Ed. (2015).

76. J. C. Ruel-Bergeron, et al., Global Update and Trends of Hidden Hunger, 1995-2011: The Hidden Hunger Index. PLoS One 10, e0143497 (2015).

77. J. Fanzo, et al., 2018 Global Nutrition Report: Shining a light to spur action on nutrition (2018) (June 12, 2020).

78. J. R. Bogard, et al., Nutrient composition of important fish species in Bangladesh and potential contribution to recommended nutrient intakes. J. Food Compost. Anal. 42, 120–133 (2015).

79. W. W. L. Cheung, et al., Shrinking of fishes exacerbates impacts of global ocean changes on marine ecosystems. Nat. Clim. Chang. 3, 254–258 (2013).

80. L. A. Rogers, et al., Shifting habitats expose fishing communities to risk under climate change. Nat. Clim. Chang. 9, 512–516 (2019).

81. R. Karimi, C. L. Folt, Beyond macronutrients: element variability and multielement stoichiometry in freshwater invertebrates. Ecol. Lett. 9, 1273–1283 (2006).

82. M. T. Arts, M. T. Brett, M. Kainz, Lipids in Aquatic Ecosystems (Springer Science & Business Media, 2009).

83. R. DeFries, et al., Global nutrition. Metrics for land-scarce agriculture. Science 349, 238–240 (2015).

84. N. Roos, M. A. Wahab, C. Chamnan, S. H. Thilsted, The Role of Fish in Food-Based Strategies to Combat Vitamin A and Mineral Deficiencies in Developing Countries. J. Nutr. 137, 1106–1109 (2007).

85. EPA, Guidance for Assessing Chemical Contaminant Data for Use in Fish Advisories (2000).

86. J. E. Byrnes, J. J. Stachowicz, The consequences of consumer diversity loss: different answers from different experimental designs. Ecology 90, 2879–2888 (2009).

87. T. V. Elzhov, K. M. Mullen, A.-N. Spiess, B. Bolker, minpack.lm: R interface to the Levenberg- Marquardt nonlinear least-squares algrothim found in MINPACK, pluss support for bounds (2013).

88. R Core Team, R: A Language and Environment for Statistical Computing (2019).

89. F. Baty, et al., A Toolbox for Nonlinear Regression in R : The Package nlstools. J. Stat. Softw. 66, 1–21 (2015).

90. Food and Agriculture Organization/ International Network of Food Data Systems (INFOODS), “FAO / INFOODS Guidelines for Checking Food Composition Data prior to the publication of a User Table / Database - Version 1.0” (2012).

91. Food and Agriculture Organization of the United Nations, “FAOSTAT. Calculated from food balance sheets” (2016).

92. T. Cashion, F. Le Manach, D. Zeller, D. Pauly, Most fish destined for fishmeal production are food- grade fish. Fish Fish 18, 837–844 (2017).

93. E. Laliberté, P. Legendre, A distance-based framework for measuring functional diversity from multiple traits. Ecology 91, 299–305 (2010).

