## Supplementary Materials for "Aquatic biodiversity enhances multiple nutritional benefits to humans"

**This PDF file includes:**

**Part A: Supplementary Figures and Tables (Figures S1-S12; Tables S1 – S14)**
**Part B: Supplementary Methods**
**Supplementary References**

**Part A: Supplementary Figures and Tables**

#### Part A: Supplementary Figures and Tables

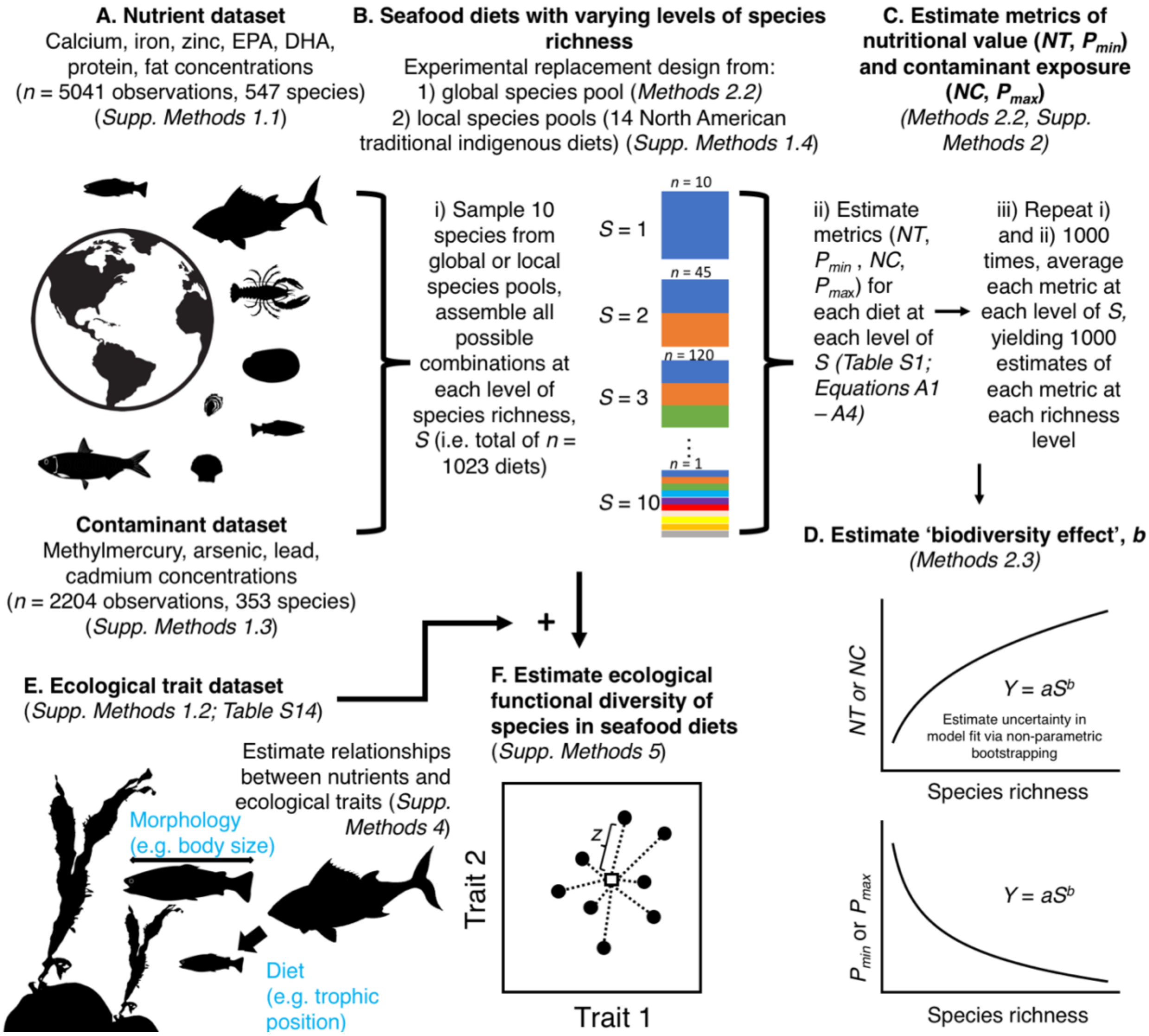

**Figure S1.** Graphical overview of some of the analyses used in this study, and references to relevant methods and definitions in the text. We began by assembling datasets of nutrient and contaminant concentrations in seafood tissues (A, Supplementary Methods 1.1). From these datasets, we assembled potential seafood diets at varying levels of species richness sourced from species pools at global and local scales, following a design analogous to a biodiversity-ecosystem function experiment with a replacement design (B, Methods 2.2, Supplementary

Methods 1.4). We then estimated metrics of nutritional value and contaminant exposure (C, Methods 2.2 and Supplementary Methods 2), and the ‘biodiversity effect’ (D, Methods 2.3). Finally, using a dataset of ecological traits for the species in seafood nutrient dataset (E, Supplementary Methods 1.2), we estimated the ecological functional diversity of seafood diets, to assess the relationship between metrics of nutritional value and ecological functional diversity (F, Supplementary Methods 5).

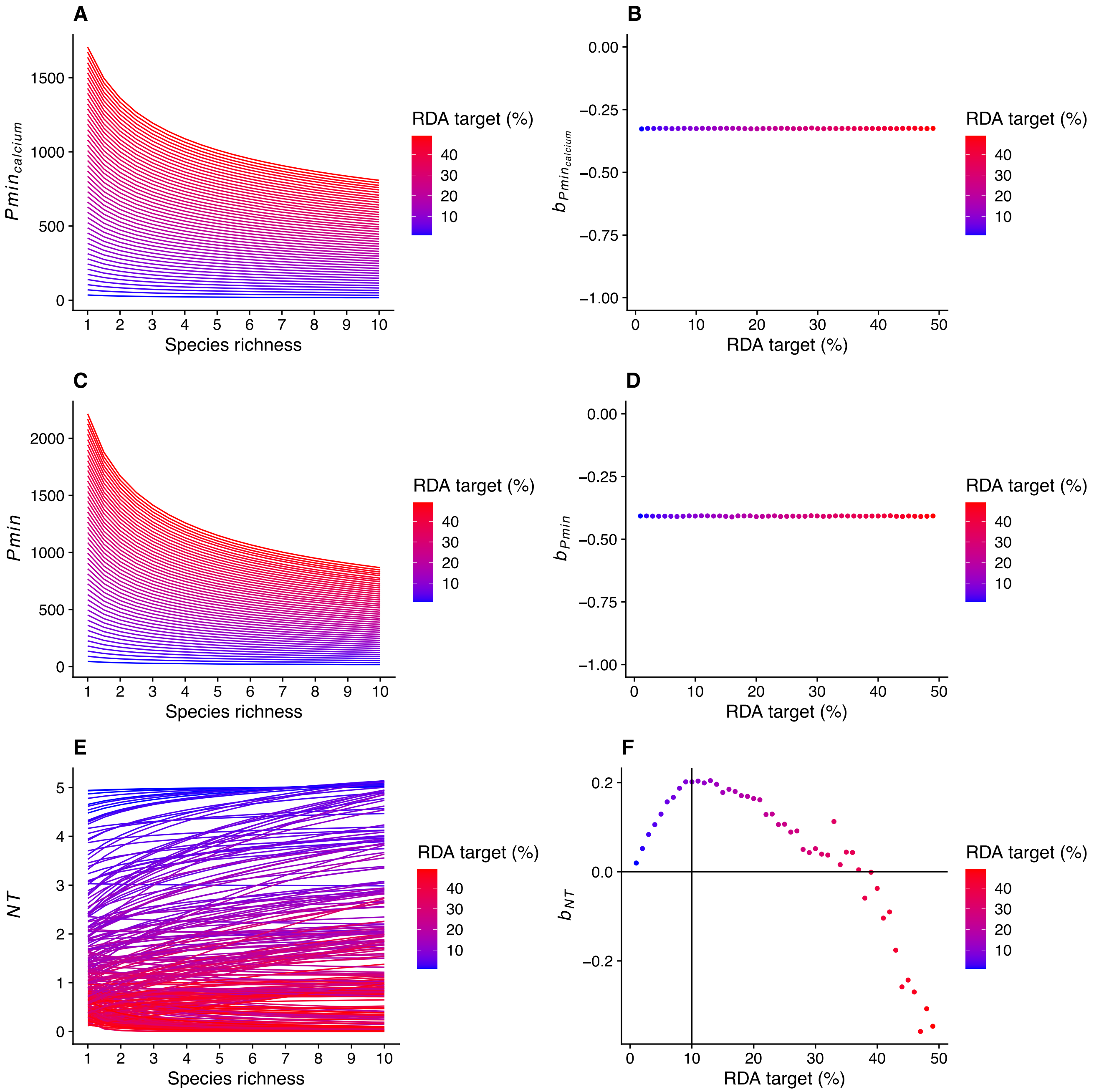

**Figure S2.** Minimum portion size ($P_{min}$, Main Text, Methods 1.1) required to reach RDA targets for a single nutrient (e.g. calcium) (**A**) and for five micronutrients (**C**) depends on species richness and the percentage of RDA considered in the target. The effect of biodiversity on $P_{min}$, (estimated as the slope parameter, $b_{Pmin}$, in Equation 3, Main Text) remains constant across all percentages of RDA, in the case of $b_{Pmin}$ for a single nutrient (e.g. calcium), (**B**) and five micronutrients simultaneously (**D**). Each line shown in **A** and **C** is the mean $P_{min}$ from 1000 simulated diets sampled from the global seafood nutrient dataset. Colours correspond to calculations of $P_{min}$ using different RDA target levels, from 1% to 50% RDA. The effect of biodiversity on the number of nutrients that reach RDA targets (*NT*, Main Text, Methods 1.1) depends on the RDA target level (**E, F**). Number of nutrients that reach RDA targets per 100g edible portion increases with seafood species richness (**E**). Colours correspond to calculations of *NT* using different RDA target levels, from 1% to 50% RDA. Each line shown here is the mean *NT* from 1000 simulated diets sampled from the global seafood nutrient dataset. The effect of biodiversity (estimated as the slope parameter, $b_{NT}$, in Equation 2, Main Text) on the number of nutrient targets reached (*NT*) is positive over a range of RDA target levels from approximately 1-40% (**F**).

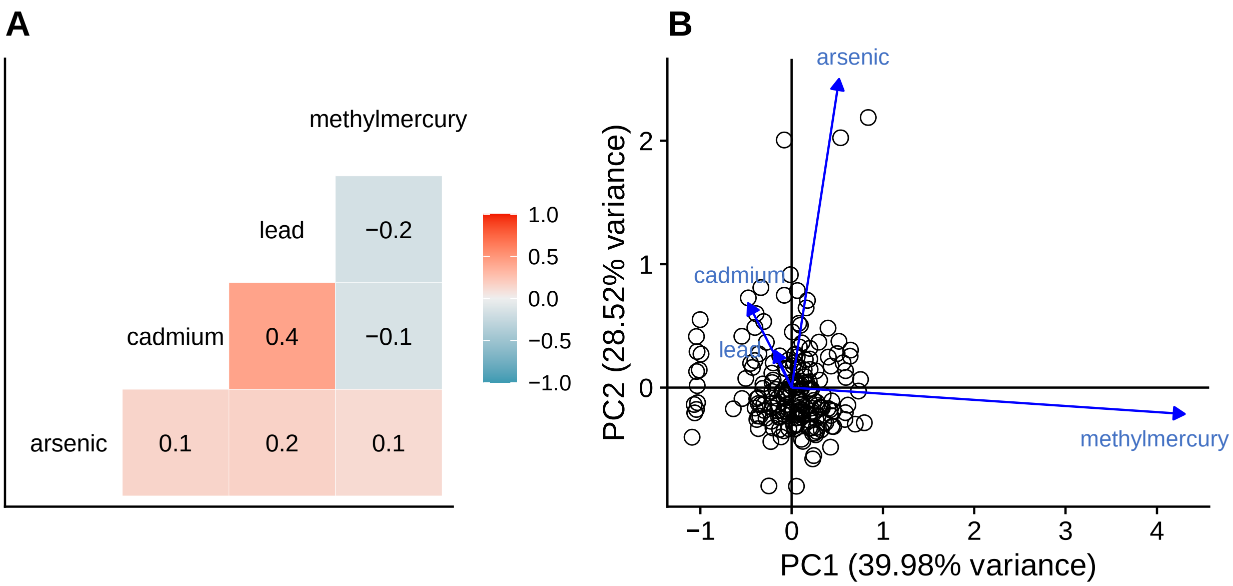

**Figure S3.** Pairwise Pearson correlation coefficients (A) and principal components analysis (B) of contaminant concentrations in muscle tissues of 200 North American seafood species (see SI Supplementary Methods 1.3 for information on contaminant data).

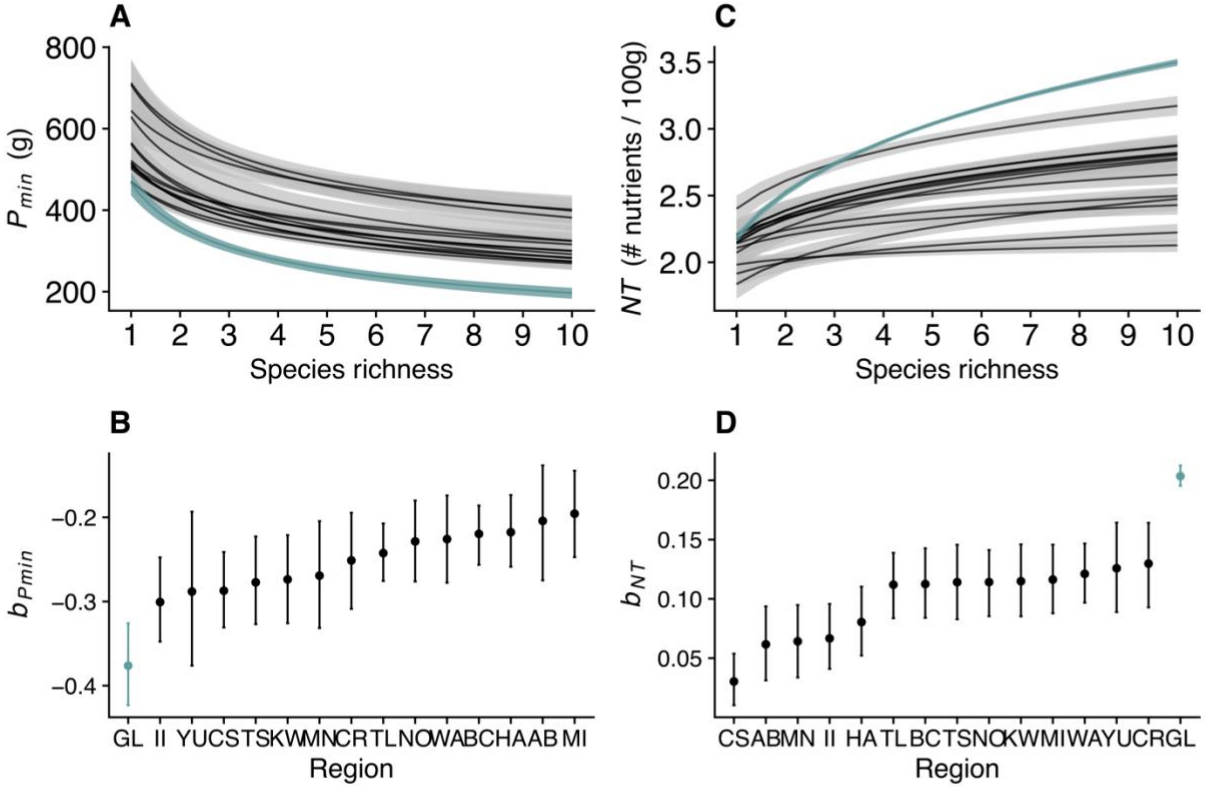

**Figure S4.** At global and local scales, biodiversity enhances nutritional benefits in terms of two metrics of nutritional benefit: minimum portion size required to reach five micronutrient targets, $P_{min}$, (A, B) and the number of nutrients that meet an RDA target, *NT* (C, D). Increasing seafood species richness sourced from local and global scales reduces the minimum portion size required to reach five micronutrient RDA targets (A, black lines are the fit of Equation 3 for each of fourteen traditional Indigenous diets in North America (local scales), green line is for a diet sourced from the global seafood market (global scale)). Increasing species richness increases *NT* at local and global scales (C, colour coding as in A). Each point in panel B and D corresponds to the *b* parameter estimate from Equation 3 (panel A) and Equation 2 (panel C) for one of fourteen local Indigenous diets (Table S12) and the global diet (GL; standardized to 40 species, Supplementary Methods 3). Points are mean ± 95% CI from non-parametric bootstrapping of the fit of Equations 2 or 3 to randomly assembled diets drawing from the species included in the

North American Indigenous diet species' lists (Supplementary Methods, section 1.4). Names of regions for each local diet are represented in two-letter abbreviations listed in Table S12.

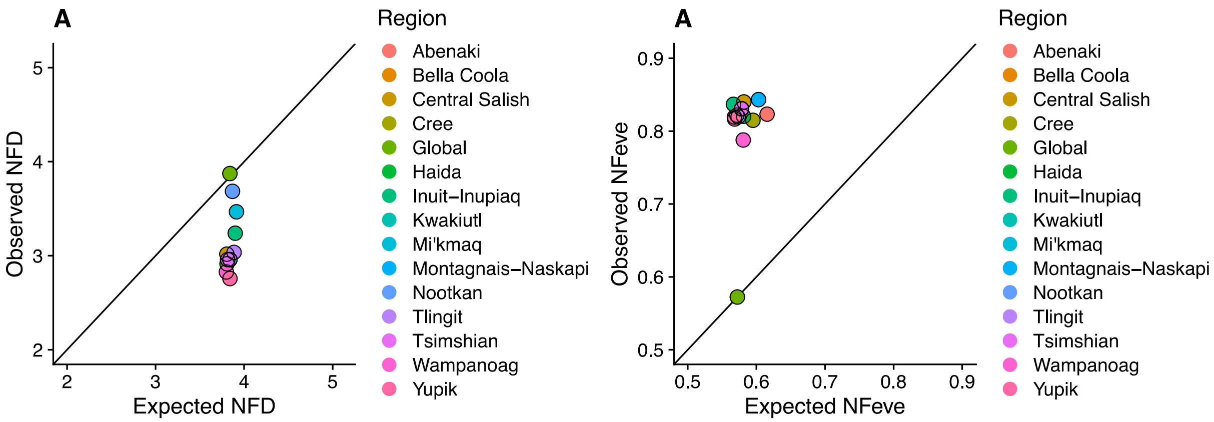

**Figure S5.** Observed *vs* expected nutritional functional diversity (*NFD*) and functional evenness (*NFEve*) in local North American traditional Indigenous seafood diets and global seafood diets. Local Indigenous diets tend to have lower *NFD* and higher *NFEve* than global seafood diets standardized to the same number of species (40 species); see Supplementary Methods section 3.

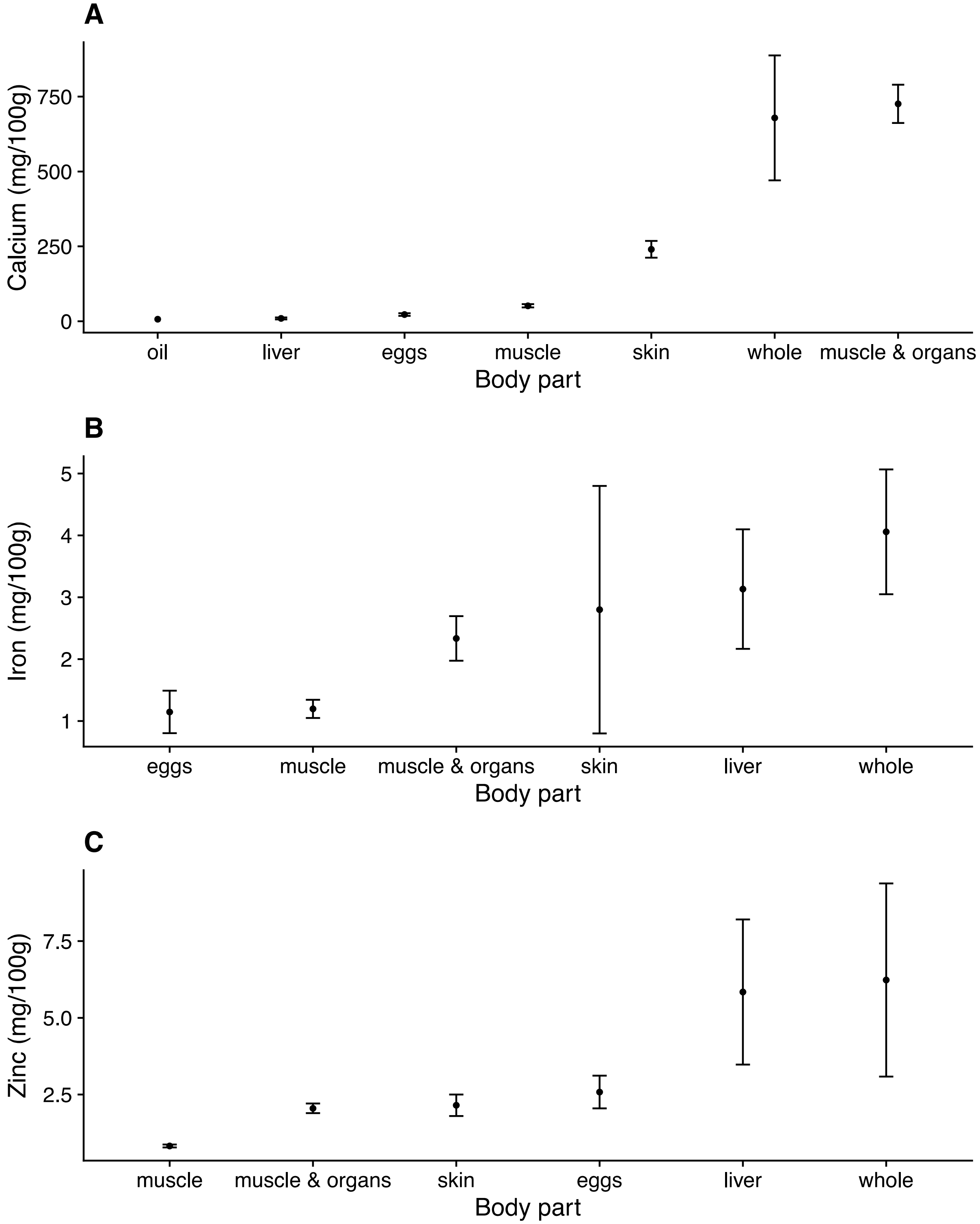

**Figure S6.** Variation in nutrient content per portion of finfish species associated with different body parts. Among finfish species, nutrient content varies by body part in the edible portion. Fish species that are eaten whole or whose edible portions include organs such as skin, liver or bones have higher nutrient content for some nutrients than those whose edible portions are restricted to muscle tissue. Note that different edible portions may come from different species, and not all tissues are available for all species. (A) calcium, $n$ = 343 observations, (ANOVA, $F_{6, 336} = 33.42$, $p < 0.01$), (B) iron, $n$ = 316 observations (ANOVA, $F_{5, 310} = 4.36$, $p < 0.01$), (C) zinc $n$ = 299 observations (ANOVA, $F_{5, 293} = 41.98$, $p < 0.01$). Points are mean ± standard error.

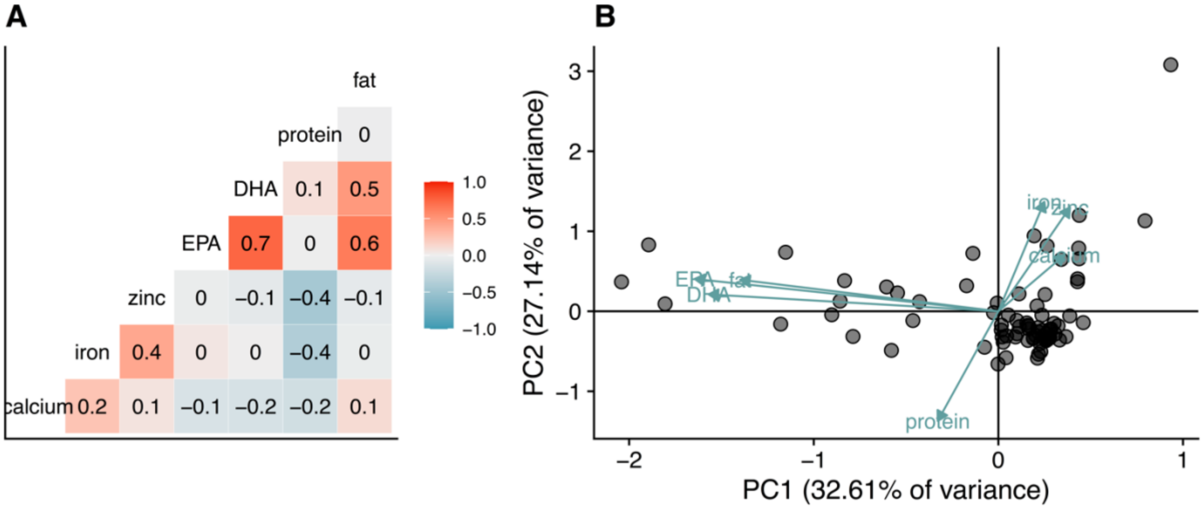

**Figure S7.** A) Pairwise Pearson correlation coefficients among concentrations of seven nutrients (five micronutrients plus protein and fat) in 120 species in the nutrient dataset for which data on all seven nutrients were available. B) Principal components analysis of seafood species' nutrient concentrations in edible portions for all seven nutrients showing trade-offs between the essential microelements (iron, calcium and zinc) and essential fatty acids (EPA and DHA).

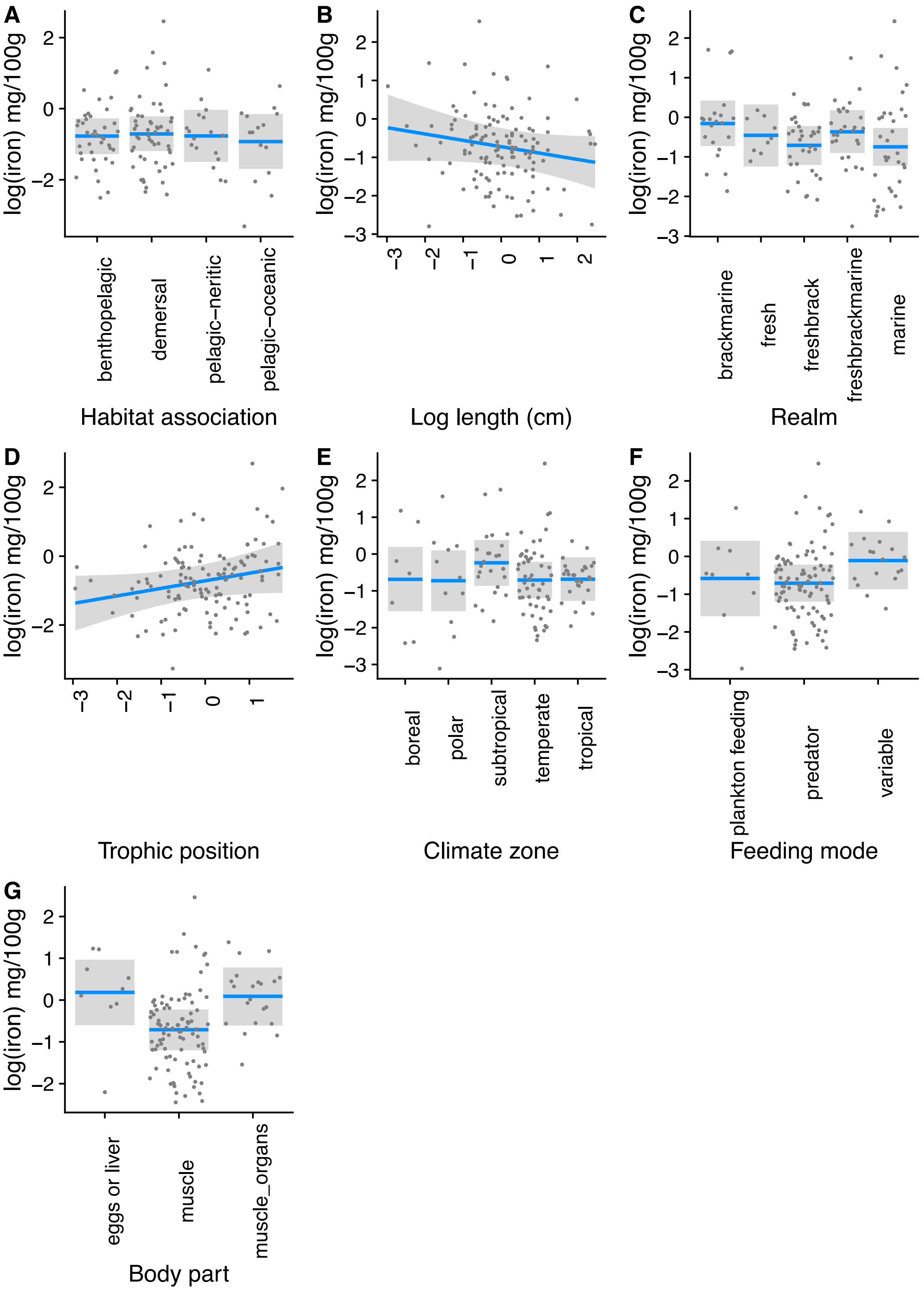

A
log(iron) mg/100g
benthopelagic
demersal
pelagic–neritic
pelagic–oceanic
Habitat association
B
log(iron) mg/100g
Log length (cm)
C
log(iron) mg/100g
brackmarine
fresh
freshbrack
freshbrackmarine
marine
Realm
D
log(iron) mg/100g
Trophic position
E
log(iron) mg/100g
boreal
polar
subtropical
temperate
tropical
Climate zone
F
log(iron) mg/100g
plankton feeding
predator
variable
Feeding mode
G
log(iron) mg/100g
eggs or liver
muscle
muscle_organs
Body part

**Figure S8.** Iron concentrations in edible finfish tissues vary with species' ecological traits. Partial regression plots from phylogenetic least squares regression (Supplementary Methods section 4) showing the effects of ecological traits on iron concentrations in seafood species edible tissues (finfish only), while holding all other variables constant ($n$ = 123 species, $R^2$ = 0.26; for full model results and regression coefficient estimates, see Table S4). Note that different edible portions may come from different species, and data for nutrients in all tissue types were not available for all species; see Table S14 for predictors.

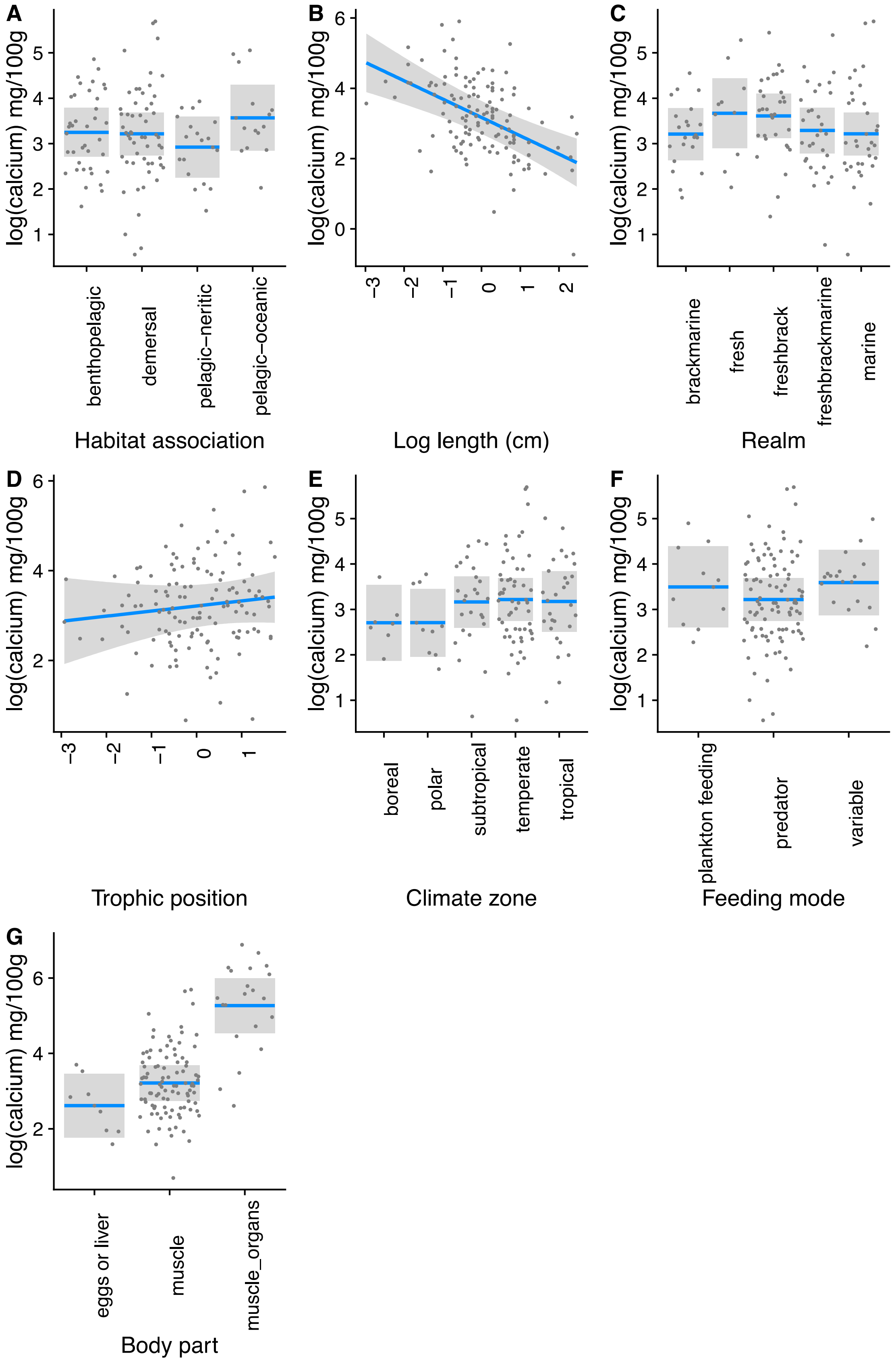
A
B
C
D
E
F
G
log(calcium) mg/100g
Habitat association
benthopelagic
demersal
pelagic–neritic
pelagic–oceanic
Log length (cm)
Realm
brackmarine
fresh
freshbrack
freshbrackmarine
marine
Trophic position
Climate zone
boreal
polar
subtropical
temperate
tropical
Feeding mode
plankton feeding
predator
variable
Body part
eggs or liver
muscle
muscle_organs

**Figure S9.** Calcium concentrations in edible finfish tissues vary with species' ecological traits. Partial regression plots from phylogenetic least squares regression (Supplementary Methods section 4) showing the effects of ecological traits on calcium concentrations in seafood species edible tissues (finfish only), while holding all other variables constant ($n$ = 127 species, $R^2$ = 0.60; for full model results and regression coefficient estimates, see Table S5). Note that different edible portions may come from different species, and data for nutrients in all tissue types were not available for all species; see Table S14 for predictors.

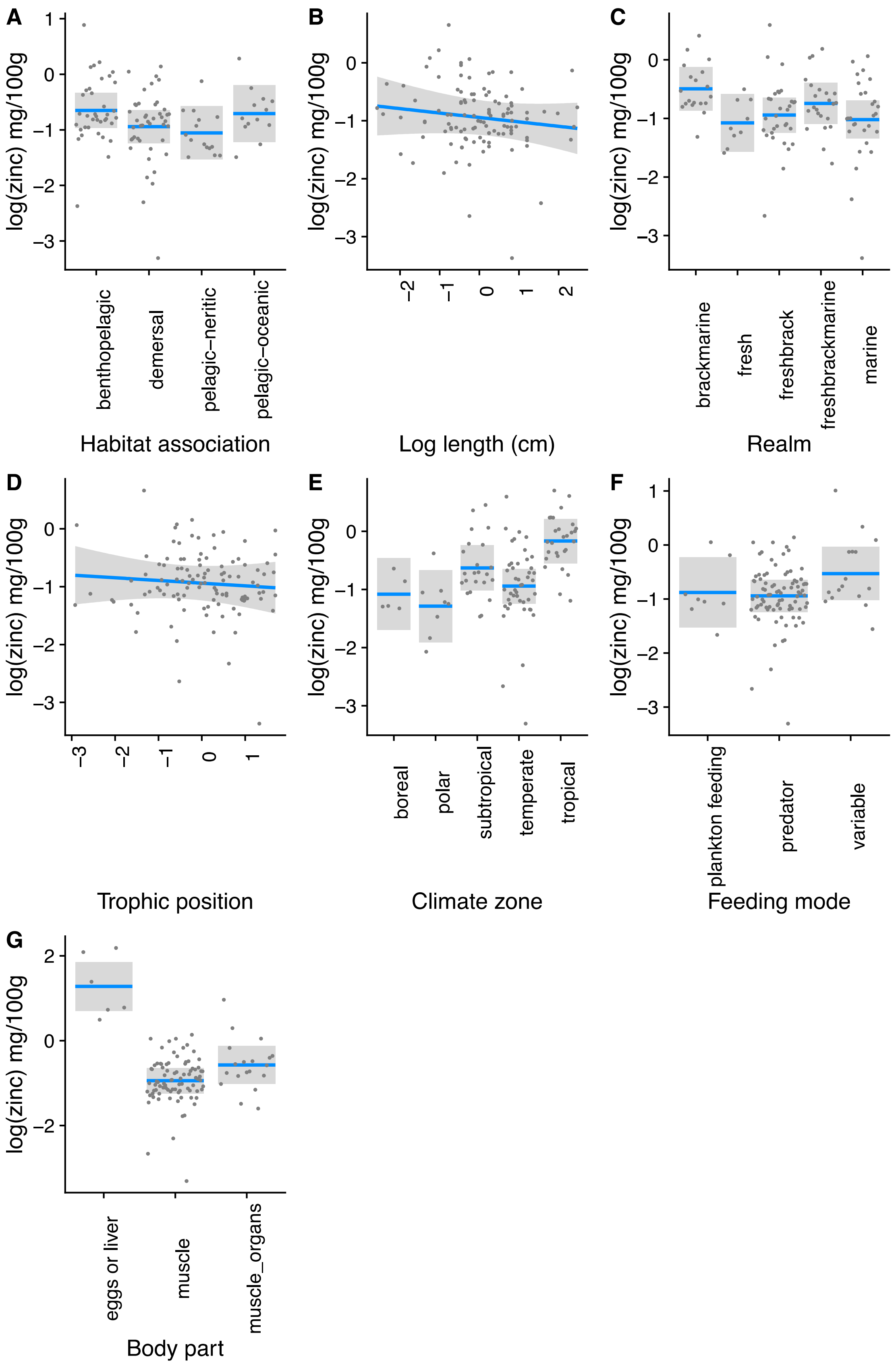
A
B
C
D
E
F
G
log(zinc) mg/100g
Habitat association
benthopelagic
demersal
pelagic–neritic
pelagic–oceanic
Log length (cm)
Realm
brackmarine
fresh
freshbrack
freshbrackmarine
marine
Trophic position
Climate zone
boreal
polar
subtropical
temperate
tropical
Feeding mode
plankton feeding
predator
variable
Body part
eggs or liver
muscle
muscle_organs

**Figure S10.** Zinc concentrations in edible finfish tissues vary with species' ecological traits. Partial regression plots from phylogenetic least squares regression (Supplementary Methods section 4) showing the effects of ecological traits on zinc concentrations in seafood species edible tissues (finfish only), while holding all other variables constant (*n* = 110 species, $R^2$ = 0.61; for full model results and regression coefficient estimates, see Table S6). Note that different edible portions may come from different species, and data for nutrients in all tissue types were not available for all species; see Table S14 for predictors.

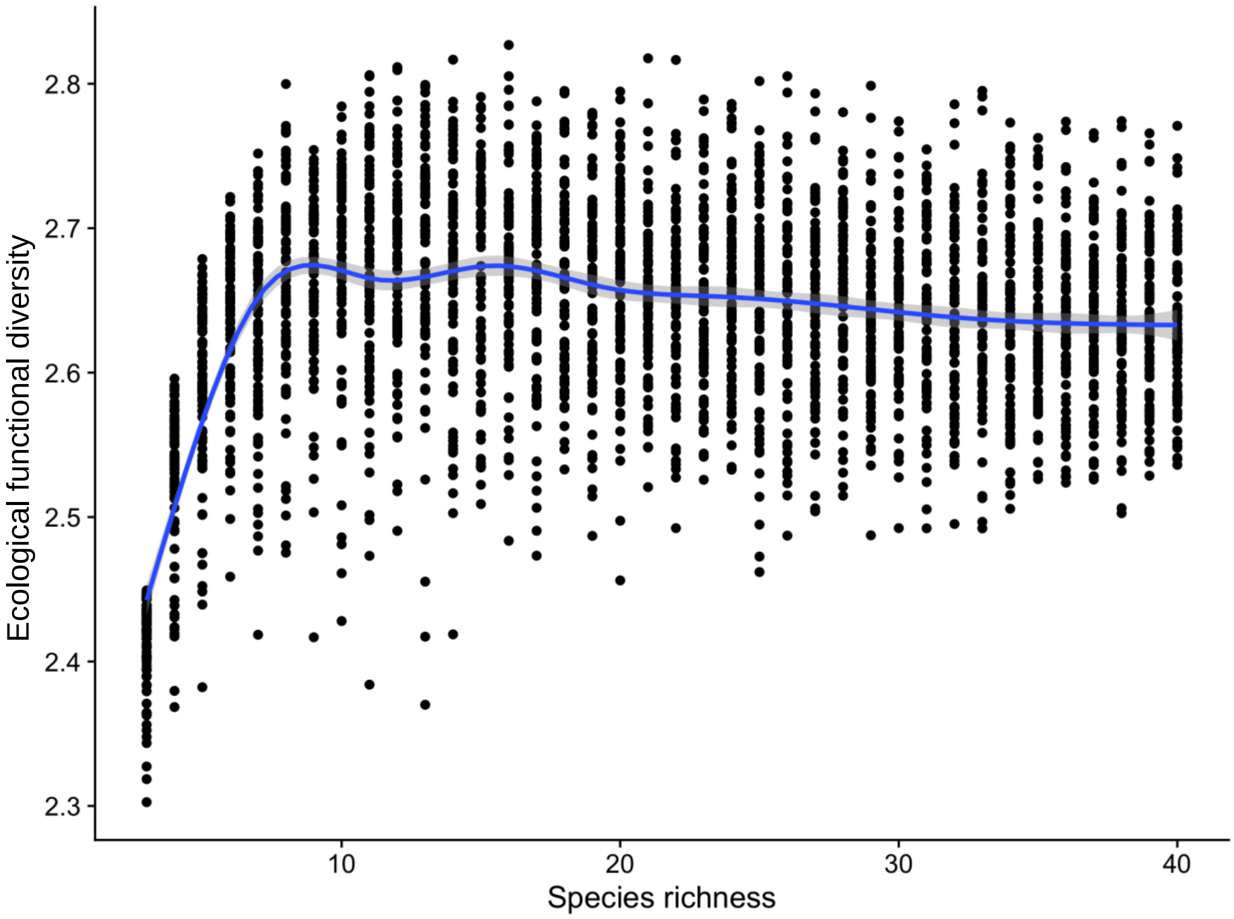

**Figure S11.** Ecological functional diversity of species from the global seafood species pool (Main Text Methods 2.3, Supplementary Methods 5) measured as functional dispersion (i.e. the mean difference in multidimensional trait space of each individual species from the centroid of all species, **Figure 6A**) increases with species richness from three to ten species. Each point represents the ecological functional diversity estimated from randomly assembled seafood diets (resampled from the global seafood species dataset 100 times) at different levels of species richness (from three species to 40 species).

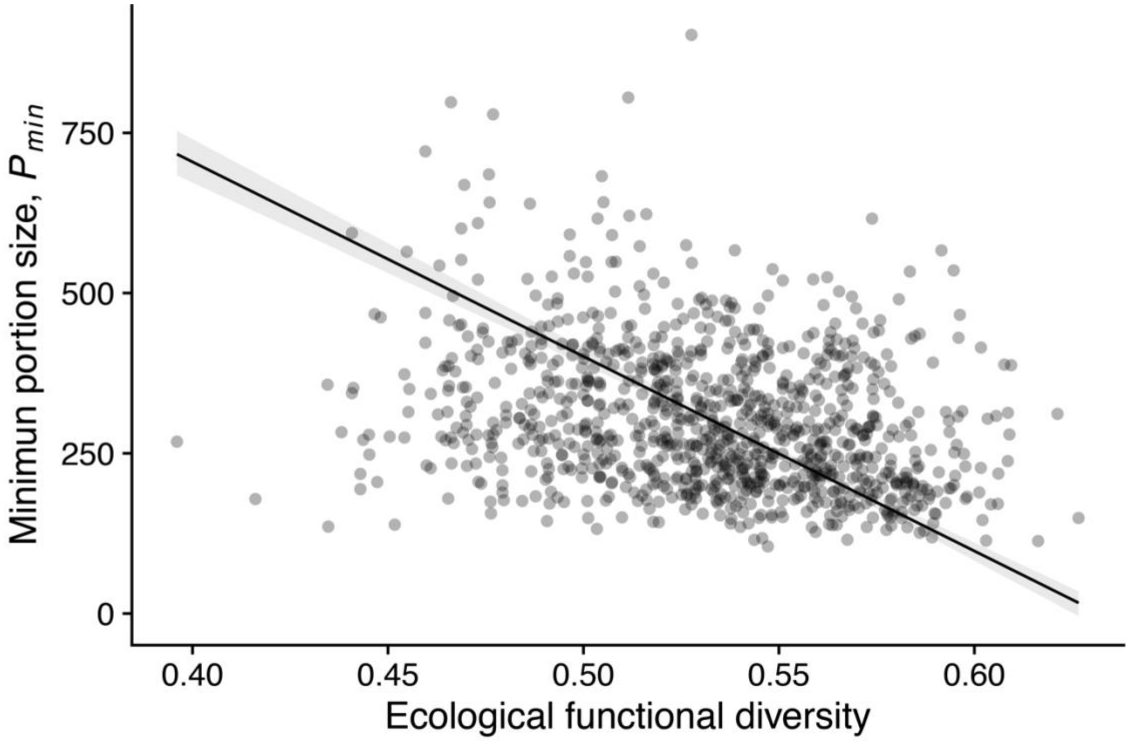

**Figure S12.** Diets with higher levels of ecological functional diversity (*EFD*, Supplementary Methods 5, Table S1) are associated with lower minimum portion sizes ($P_{min}$) required to reach five micronutrient RDA targets simultaneously. Each point corresponds to a seafood species diet with ten species (i.e. richness, $S = 10$), and 1000 randomly assembled diets from the global seafood species dataset are shown. For each of the 1000 simulated diets, $P_{min}$ and *EFD* were calculated (Methods 1.3 and Supplementary Methods 5). Reduced major axis regression slope =-349.92, 95% CI -1436.16, -1268.86, $R^2 = 0.0048$.

**Table S1. Definitions of key terms and references.**

| Term | Definition |
|---|---|
| **Minimum portion size, $P_{min}$** | Minimum amount of edible seafood tissue (g) required to reach a given RDA target for one nutrient or a set of nutrients.<br><br>Given the total number of species *S* indexed by *j* and total number of nutrients *N* indexed by *i*,<br><br>$$P_{min} = \arg\min_{P}\left(\sum_{i=1}^{N} \mathbb{1}\left\{\sum_{j=1}^{S} \frac{P}{S} content_{ij} \geq RDAtarget_i\right\} = N\right) \quad (A1)$$<br><br>where $P$ is the total amount of edible seafood tissue; $content_{ij}$ is the nutritional content (mg) per 100g portion of species *j* and nutrient *i*, and $RDAtarget_i$ is the RDA target for nutrient *i*. The $\mathbb{1}$ refers to an indicator function, which takes on the value of 1 when the expression in the curly brackets is true (i.e. when the total nutrient content in the diet of *S* species, shown on the left-hand side of the "≥" is greater than or equal to the right-hand side of the expression, the RDA target), and 0 when the expression in the curly brackets is false (i.e. when total nutrient content summed across *S* species is less than the RDA target). The $\arg\min_{P}$ denotes that *P* is the argument that is being minimized. $P_{min}$ is the minimum value of *P* (amount of edible tissue) required to reach a single RDA target (when *N* = 1) or set of RDA targets (when *N* > 1). In this paper, we considered the case when *N* = 1 for one nutrient, and *N* = 5 for five nutrients simultaneously. In other words, $P_{min}$ is the minimum amount of edible tissue that reaches or exceeds a given number of RDA targets. |
| **Number of nutrients, *NT*** | Number of nutrients that reach a specific RDA target (for example, 10% RDA) in a single 100g portion of seafood (here *NT* ranges from 0-5).<br><br>Given the total number of species *S* indexed by *j* and total number of nutrients (calcium, iron, zinc, EPA, DHA) *N* indexed by *i*, |

| | |
|---|---|
| | $$NT = \sum_{i=1}^{N} \mathbb{1}\left\{\sum_{j=1}^{S} \frac{1}{S} content_{ij} \geq RDAtarget_i\right\} \qquad (A2)$$ where $content_{ij}$ is the nutrient content (mg) per 100g portion of species $j$ and nutrient $i$, and $RDAtarget_i$ is the RDA target for nutrient $i$. The $\mathbb{1}$ refers to an indicator function, which takes on the value of 1 when the expression in the curly brackets is true (i.e. when the nutrient content of a 100g edible portion composed of *S* species is equal to or greater than the RDA for nutrient *i*), and 0 when the expression in the curly brackets is false (i.e. when the nutrient content of a 100g edible portion composed of *S* species is less than the RDA for nutrient *i*). |
| **Maximum portion size, $P_{max}$** | Maximum amount of edible seafood tissue (g) that may be consumed per day before one or more Provisional Tolerable Daily Intakes is exceeded. Given the total number of species, $S$, indexed by $j$ and total number of contaminants (methylmercury, arsenic, cadmium or lead), $C$, indexed by $k$, $$P_{max} = \arg\max_{P}\left(\sum_{k=1}^{C} \mathbb{1}\left\{\sum_{j=1}^{S} \frac{P}{S} content_{kj} \leq PTDI_k\right\} = C\right) \qquad (A3)$$ where $content_{kj}$ is the contaminant content (ug) per 100g of species $j$ and contaminant $k$, and $PTDI_k$ is the Provisional Tolerable Daily Intake for contaminant $k$. The $\mathbb{1}$ refers to an indicator function, which takes on the value of 1 when the expression in the curly brackets is true (i.e. when the total contaminant content in the diet of *S* species, shown on the left-hand side of the "≤" is less than or equal to the right-hand side of the expression, the PTDI), and 0 when the expression in the curly brackets is false (i.e. when total contaminant content summed across *S* species is greater than the PTDI). The $\arg\max_{P}$ denotes that *P* is the argument that is being maximized. $P_{max}$ is the maximum value of *P* (amount of edible tissue) which does not exceed one or more PTDIs. In this paper, we considered the case when *C* = 1. In other words, $P_{max}$ is the maximum amount of edible tissue that does not exceed a given PTDI (i.e. remains below the PTDI). |

| | |
|---|---|
| **Number of contaminants, *NC*** | Number of contaminants that exceed the Provisional Tolerable Daily Intake in a single 100g portion of seafood (here *NC* ranges from 0-4).<br><br>Given the total number of species *S* indexed by *j* and total number of contaminants (methylmercury, arsenic, cadmium, lead), *C*, indexed by *k*,<br><br>$$NC = \sum_{k=1}^{C} \mathbb{1}\left\{\sum_{j=1}^{S} \frac{1}{S} content_{kj} \geq PTDI_k\right\} \quad (A4)$$<br><br>where $content_{kj}$ is the contaminant content (ug) per 100g portion of species *j* and contaminant $k$, and $PTDI_k$ is the Provisional Tolerable Daily Intake for contaminant $k$. The $\mathbb{1}$ refers to an indicator function, which takes on the value of 1 when the expression in the curly brackets is true (i.e. when contaminant content of a 100g edible portion composed of *S* species is equal to or greater than the PTDI for contaminant *k*), and 0 when the expression in the curly brackets is false (i.e. when the contaminant content of a 100g edible portion composed of *S* species is less than the PTDI for contaminant *k*). |
| **Recommended dietary allowance (RDA)** | Nutritional guideline describing the average daily level of intake sufficient to meet the nutrient requirements of nearly all (97%-98%) healthy people (here we refer to women aged 19-50)(1). RDAs are measured in mg/day. |
| **RDA target** | An RDA target is met if a given edible portion of seafood meets or exceeds the RDA (or a specified portion of RDA) for a given nutrient. In this study, we considered a threshold of 10% of RDA, meaning an RDA target is 10% of RDA (mg/day). |
| **Provisional tolerable intake (weekly or daily)** | An estimate of the amount per unit body weight of a potentially harmful substance or contaminant in food or water that can be ingested over a lifetime without risk of adverse health effects (2). We used an estimate of Provisional Tolerable Daily Intake (**PTDI**) by dividing the Provisional Tolerable Weekly Intake established by Joint FAO/WHO Expert Committee on Food Additives (JECFA) by (7). |
| **Seafood** | Freshwater or marine finfish or invertebrates potentially consumed in the human diet. |

| | |
|---|---|
| **Species richness, *S*** | Number of seafood species. |
| **Ecological functional diversity, *EFD*** | The diversity of ecological traits within a community or set of species (3–5). In this study, we measured ecological functional diversity as functional dispersion following Laliberte and Legendre 2010, (4), which measures the mean distance, in multivariate trait space, of each individual species from the centroid of all species (Methods 2.3, Supplementary Methods 5). |
| **Nutritional functional diversity, *NFD*** | Estimates the diversity of nutrient concentrations in a diet based on an assessment of the entire nutritional diversity of a group represented as a functional dendrogram. Quantifies the degree of nutritional distinctiveness among species based on the dendrogram (6) (Supplementary Methods 3). |
| **Multi-nutrient profile** | The concentration of five micronutrients - calcium, iron, zinc, EPA and DHA in edible seafood tissues, characterized in multivariate space. |

**Table S2. Recommended Dietary Allowances (RDA) and Provisional Tolerable Intakes (PTWI).**

**A. Nutrients**

| Nutrient | RDA (per day) | unit | RDA target (10% RDA) |
|---|---|---|---|
| Zinc | 8 | mg | 0.8 |
| Calcium | 1000 | mg | 100 |
| Iron | 18 | mg | 1.8 |
| EPA | 1 | g | 0.1 |
| DHA | 1 | g | 0.1 |
| Protein | 46 | g | 4.6 |
| Fat | 70 | g | 7 |

**B. Contaminants**

| Contaminant | PTWI (ug/kg body weight/wk) | PTDI (ug per 70kg person per day) |
|---|---|---|
| Arsenic (total) | 15 | 150 |
| Cadmium | 7 | 70 |
| Lead | 25 | 250 |
| Methylmercury | 1.6 | 16 |

**Table S2. A)** Recommended dietary allowances (RDA) for micro- and macronutrients considered in this study. RDAs are average daily dietary intake levels that are sufficient to meet the nutrient requirement of nearly all (97 to 98%) healthy individuals in a particular life stage and gender group. Here we used the RDAs for females aged 19-50 (1, 7, 8). **B**) Provisional Tolerable Weekly Intake (PTWI) limits for heavy metals. PTWIs estimate the amount per unit body weight of a potentially harmful substance or contaminant in food or water that can be ingested over a lifetime without risk of adverse health effects. Here we used the PTWI established by the Joint FAO/WHO Expert Committee on Food Additives (JECFA)(2). To make our contaminants analysis consistent with the nutrient analyses, which was conducted for a daily diet (i.e. with reference to daily RDA) we calculated a Provisional Tolerable Daily Intake PTDI, by dividing the PTWI by 7, for a 70kg person, which allowed us to determine the upper tolerable limit per person, per day.

**Table S3. Species in the global seafood species nutrient dataset that reach RDA targets.**

| Nutrient | Crustacean % | Crustacean n | Finfish % | Finfish n | Mollusc % | Mollllusc n | All % | All n |
|---|---|---|---|---|---|---|---|---|
| Calcium | 16.67 | 18 | 32.6 | 181 | 26.32 | 38 | 30.38 | 237 |
| DHA | 44 | 25 | 72.77 | 224 | 60 | 30 | 68.82 | 279 |
| EPA | 50 | 26 | 46.82 | 220 | 66.67 | 30 | 49.28 | 276 |
| Fat | 9.52 | 42 | 16.08 | 398 | | | 15.45 | 440 |
| Iron | 47.06 | 17 | 23.03 | 178 | 82.5 | 40 | 34.89 | 235 |
| Protein | 100 | 33 | 100 | 322 | 100 | 60 | 100.00 | 415 |
| Zinc | 100 | 10 | 46.34 | 164 | 90.91 | 33 | 56.04 | 207 |

**Table S3.** Percentage of species in the seafood nutrient dataset that reach 10% of RDA in a 100g edible portion, and total number of species (*n*) grouped by taxonomic group.

**Table S4. Dependence of Iron (Fe) concentrations in edible finfish tissues on ecological traits.**

| Term | $w_{ip}$ | 1 | 2 | 3 | 4 | 5 | 6 | 7 | 8 | $\beta_i$ | Lower 95% CI | Upper 95% CI |
|---|---|---|---|---|---|---|---|---|---|---|---|---|
| Body part: muscle | 1 | + | + | + | + | + | + | + | + | **-0.75** | **-1.45** | **-0.05** |
| Body part: muscle & organs (incl. bones) | 1 | + | + | + | + | + | + | + | + | 0.04 | -0.77 | 0.86 |
| Length | 0.76 | -0.21 | | -0.13 | -0.16 | -0.13 | -0.19 | | | -0.19 | -0.41 | 0.02 |
| Trophic position | 0.8 | 0.14 | 0.17 | | | | 0.15 | 0.08 | 0.19 | 0.15 | -0.06 | 0.35 |
| Feeding mode: hunting | 0.35 | | + | + | | | | | + | -0.26 | -1.08 | 0.57 |
| Feeding mode: variable | 0.35 | | + | + | | | | | + | 0.25 | -0.6 | 1.1 |
| Climate zone: polar | 0.05 | | | | + | | | | | -0.2 | -1.19 | 0.8 |
| Climate zone: subtropical | 0.05 | | | | + | | | | | 0.3 | -0.55 | 1.16 |
| Climate zone: temperate | 0.05 | | | | + | | | | | -0.18 | -0.98 | 0.62 |
| Climate zone: tropical | 0.05 | | | | + | | | | | -0.04 | -0.9 | 0.82 |
| Diet source: demersal | 0.08 | | | | | + | + | + | + | 0.03 | -0.4 | 0.46 |
| Diet source: pelagic-neritic | 0.08 | | | | | + | + | + | + | 0.24 | -0.35 | 0.84 |
| Diet source: oceanic | 0.08 | | | | | + | + | + | + | -0.09 | -0.77 | 0.59 |
| Realm | | | | | | | | | | | | |
| Pagel's $\lambda$ | | 0 | 0 | 0 | 0 | 0 | 0 | 0 | 0 | | | |
| $R^2$ | | 0.16 | 0.16 | 0.16 | 0.17 | 0.15 | 0.16 | 0.14 | 0.17 | | | |
| Δ AICc | | 0 | 1.86 | 2.83 | 4.9 | 5.88 | 6.04 | 7.07 | 7.16 | | | |
| Akaike weight | | 0.5 | 0.2 | 0.12 | 0.04 | 0.03 | 0.02 | 0.01 | 0.01 | | | |
| Cumulative weight | | 0.5 | 0.7 | 0.82 | 0.86 | 0.89 | 0.91 | 0.93 | 0.94 | | | |

**Table S4.** Results from phylogenetic least squares regression model selection and model averaging for models relating iron concentrations in edible finfish species tissues to finfish species' traits (*n* = 123 species) (see Supplementary Methods 4 for full model description). Models are ranked in order of increasing AICc differences (Δ AICc). All models in the 95%

confidence model set are shown here (cumulative sum of Akaike weights <= 0.95). Akaike weights indicate the relative likelihood of a model, given the set of models being considered. Model-averaged standardized regression coefficients, $\beta_i$, with 95% CI, were averaged across all models in the best set and weighted by each model's Akaike weight. $\beta_i$ were calculated with $\beta_i$ = 0 when variable *i* was not included in a model. $\beta_i$ with 95% confidence intervals not encompassing 0 are shown in bold. Relative variable importance ($w_{ip}$) is the sum of the relative variable importance across all models including that variable (*10*). Traits for which there is not a coefficient estimate did not appear in any of the models in the top 95% set (with cumulative Akaike weight <= 0.95). Note that different edible portions may come from different species, and not all tissues are available for all species.

**Table S5. Dependence of Calcium (Ca) concentrations in edible finfish tissues on ecological traits.**

| Term | $w_{ip}$ | 1 | 2 | 3 | 4 | $\beta_i$ | Lower 95% CI | Upper 95% CI |
|---|---|---|---|---|---|---|---|---|
| Body part: muscle | 1.00 | + | + | + | + | 0.64 | -0.05 | 1.33 |
| Body part: muscle & organs (incl. bones) | 1.00 | + | + | + | + | 2.83 | 2.04 | 3.63 |
| Length | 1.00 | -0.45 | -0.45 | -0.51 | -0.50 | **-0.47** | **-0.68** | **-0.26** |
| Trophic position | 0.45 | -0.04 | | | -0.05 | -0.04 | -0.24 | 0.16 |
| Feeding mode: hunting | 0.29 | | + | | | 0.01 | -0.67 | 0.68 |
| Feeding mode: variable | 0.29 | | + | | | 0.36 | -0.41 | 1.13 |
| Diet source: demersal | 0.35 | | | + | + | -0.18 | -0.59 | 0.24 |
| Diet source: pelagic-neritic | 0.35 | | | + | + | -0.51 | -1.06 | 0.05 |
| Diet source: pelagic-oceanic | 0.35 | | | + | + | 0.06 | -0.56 | 0.69 |
| Climate zone | | | | | | | | |
| Realm | | | | | | | | |
| Pagel's $\lambda$ | | 0 | 0 | 0 | 0 | | | |
| $R^2$ | | 0.56 | 0.56 | 0.57 | 0.57 | | | |
| Δ AICc | | 0.00 | 0.42 | 0.66 | 2.76 | | | |
| Akaike weight | | 0.34 | 0.28 | 0.24 | 0.09 | | | |
| Cumulative weight | | 0.34 | 0.61 | 0.86 | 0.94 | | | |

**Table S5.** Results from phylogenetic least squares regression model selection and model averaging for models relating calcium concentrations in edible finfish species tissues to finfish species' traits (*n* = 127 species) (see Supplementary Methods 4 for full model description). Models are ranked in order of increasing AICc differences (Δ AICc). All models in the 95% confidence model set are shown here (cumulative sum of Akaike weights <= 0.95). Akaike weights indicate the relative likelihood of a model, given the set of models being considered. Model-averaged standardized regression coefficients, $\beta_i$, with 95% CI, were averaged across all models in the best set and weighted by each model's Akaike weight. $\beta_i$ were calculated with $\beta_i$ = 0 when variable *i* was not included in a model. $\beta_i$ with 95% confidence intervals not encompassing 0 are shown in bold. Relative variable importance ($w_{ip}$) is the sum of the relative variable importance across all models including that variable (*10*). Traits for which there is not a coefficient estimate did not appear in any of the models in the top 95% set (with cumulative Akaike weight <= 0.95). Note that different edible portions may come from different species, and not all tissues are available for all species.

**Table S6. Dependence of Zinc (Zn) concentrations in edible finfish tissues on ecological traits.**

| Term | $w_{ip}$ | 1 | 2 | $\beta_i$ | Lower 95% CI | Upper 95% CI |
|---|---|---|---|---|---|---|
| Body part: muscle | 1.00 | + | + | **-2.27** | **-2.84** | **-1.70** |
| Body part: muscle & organs (incl. bones) | 1.00 | + | + | **-1.92** | **-2.61** | **-1.24** |
| Feeding mode: hunting | 0.91 | + | | -0.37 | -0.83 | 0.08 |
| Feeding mode: variable | 0.91 | + | | 0.17 | -0.38 | 0.73 |
| Climate zone: polar | 1.00 | + | + | -0.10 | -0.83 | 0.64 |
| Climate zone: subtropical | 1.00 | + | + | 0.50 | -0.11 | 1.11 |
| Climate zone: temperate | 1.00 | + | + | 0.17 | -0.41 | 0.76 |
| Climate zone: tropical | 1.00 | + | + | **1.01** | **0.37** | **1.64** |
| Trophic position | 0.09 | | -0.13 | **-0.13** | **-0.26** | **-0.01** |
| Diet source | | | | | | |
| Length | | | | | | |
| Realm | | | | | | |
| Pagel's $\lambda$ | | 0 | 0 | | | |
| $R^2$ | | 0.56 | 0.53 | | | |
| Δ AICc | | 0.00 | 4.57 | | | |
| Akaike weight | | 0.81 | 0.08 | | | |
| Cumulative weight | | 0.81 | 0.90 | | | |

**Table S6.** Results from phylogenetic least squares regression model selection and model averaging for models relating zinc concentrations in edible finfish species tissues to finfish species' traits ($n$ = 113 species) (see Supplementary Methods 4 for full model description). Models are ranked in order of increasing AICc differences (Δ AICc). All models in the 95% confidence model set are shown here (cumulative sum of Akaike weights <= 0.95). Akaike weights indicate the relative likelihood of a model, given the set of models being considered. Model-averaged standardized regression coefficients, $\beta_i$, with 95% CI, were averaged across all models in the best set and weighted by each model's Akaike weight. $\beta_i$ were calculated with $\beta_i$ =

0 when variable *i* was not included in a model. $\beta_i$ with 95% confidence intervals not encompassing 0 are shown in bold. Relative variable importance ($w_{ip}$) is the sum of the relative variable importance across all models including that variable (*10*). Traits for which there is not a coefficient estimate did not appear in any of the models in the top 95% set (with cumulative Akaike weight <= 0.95). Note that different edible portions may come from different species, and not all tissues are available for all species.

**Table S7. Dependence of DHA concentrations in edible finfish tissues on ecological traits.**

| Term | $w_{ip}$ | 1 | 2 | $\beta_i$ | Lower 95% CI | Upper 95% CI |
|---|---|---|---|---|---|---|
| Body part: muscle | 1.00 | + | + | **-2.04** | **-3.01** | **-1.07** |
| Body part: muscle & organs (incl. bones) | 1.00 | + | + | **-1.45** | **-2.59** | **-0.30** |
| Feeding mode: hunting | 1.00 | + | + | -0.33 | -0.96 | 0.30 |
| Feeding mode: variable | 1.00 | + | + | -0.45 | -1.18 | 0.27 |
| Length | 1.00 | 0.03 | -0.03 | 0.00 | -0.21 | 0.21 |
| Diet source: demersal | 0.53 | + | | -0.01 | -0.38 | 0.37 |
| Diet source: pelagic-neritic | 0.53 | + | | **0.66** | **0.14** | **1.18** |
| Diet source: oceanic | 0.53 | + | | 0.24 | -0.35 | 0.82 |
| Trophic position | 1.00 | 0.04 | 0.08 | 0.06 | -0.17 | 0.28 |
| Realm: freshwater | 1.00 | + | + | **-0.95** | **-1.76** | **-0.15** |
| Realm: freshwater or brackish | 1.00 | + | + | **-1.32** | **-2.02** | **-0.62** |
| Realm: freshwater or brackish or marine | 1.00 | + | + | -0.12 | -0.71 | 0.47 |
| Realm: marine | 1.00 | + | + | 0.03 | -0.35 | 0.41 |
| Climate zone: subtropical | 1.00 | + | + | 0.27 | -0.29 | 0.82 |
| Climate zone: temperate | 1.00 | + | + | 0.36 | -0.19 | 0.90 |
| Climate zone: tropical | 1.00 | + | + | -0.09 | -0.84 | 0.65 |
| Pagel's $\lambda$ | | 0.32 | 0.53 | | | |
| $R^2$ | | 0.42 | 0.43 | | | |
| $\Delta$ AICc | | 0.00 | 0.23 | | | |
| Akaike weight | | 0.50 | 0.45 | | | |
| Cumulative weight | | 0.50 | 0.95 | | | |

**Table S7.** Results from phylogenetic least squares regression model selection and model averaging for models relating DHA concentrations in edible finfish species tissues to finfish species' traits ($n$ = 170 species) (see Supplementary Methods 4 for full model description). Models are ranked in order of increasing AICc differences (Δ AICc). All models in the 95% confidence model set are shown here (cumulative sum of Akaike weights <= 0.95). Akaike weights indicate the relative likelihood of a model, given the set of models being considered. Model-averaged standardized regression coefficients, $\beta_i$, with 95% CI, were averaged across all models in the best set and weighted by each model's Akaike weight. $\beta_i$ were calculated with $\beta_i$ = 0 when variable *i* was not included in a model. $\beta_i$ with 95% confidence intervals not encompassing 0 are shown in bold. Relative variable importance ($w_{ip}$) is the sum of the relative variable importance across all models including that variable (*10*). Traits for which there is not a coefficient estimate did not appear in any of the models in the top 95% set (with cumulative Akaike weight <= 0.95). Note that different edible portions may come from different species, and not all tissues are available for all species.

**Table S8. Dependence of EPA concentrations in edible finfish tissues on ecological traits.**

| Term | $w_{ip}$ | 1 | 2 | $\beta_i$ | Lower 95% CI | Upper 95% CI |
|---|---|---|---|---|---|---|
| Body part: muscle | 1.00 | + | + | **-2.48** | **-3.56** | **-1.39** |
| Body part: muscle & organs (incl. bones) | 1.00 | + | + | **-1.45** | **-2.76** | **-0.14** |
| Feeding mode: plankton filtering | 1.00 | + | + | -0.51 | -2.68 | 1.67 |
| Feeding mode: hunting | 1.00 | + | + | -0.97 | -3.15 | 1.21 |
| Feeding mode: variable | 1.00 | + | + | -1.02 | -3.17 | 1.12 |
| Length | 1.00 | 0.03 | 0.10 | 0.06 | -0.18 | 0.29 |
| Climate zone: polar | 1.00 | + | + | 0.04 | -1.12 | 1.20 |
| Climate zone: subtropical | 1.00 | + | + | -0.18 | -1.27 | 0.91 |
| Climate zone: temperate | 1.00 | + | + | 0.16 | -0.90 | 1.22 |
| Climate zone: tropical | 1.00 | + | + | -0.10 | -1.32 | 1.12 |
| Trophic position | 1.00 | -0.01 | -0.07 | -0.03 | -0.29 | 0.22 |
| Realm: freshwater | 1.00 | + | + | **-1.76** | **-2.65** | **-0.87** |
| Realm: freshwater or brackish | 1.00 | + | + | **-1.64** | **-2.42** | **-0.87** |
| Realm: freshwater or brackish or marine | 1.00 | + | + | 0.06 | -0.57 | 0.70 |
| Realm: marine | 1.00 | + | + | -0.21 | -0.64 | 0.21 |
| Diet source: demersal | 0.33 | | + | 0.19 | -0.25 | 0.62 |
| Diet source: pelagic-neritic | 0.33 | | + | **0.71** | **0.12** | **1.30** |
| Diet source: oceanic | 0.33 | | + | 0.28 | -0.38 | 0.95 |
| Pagel's $\lambda$ | | 0.43 | 0.43 | | | |
| $R^2$ | | 0.46 | 0.48 | | | |
| $\Delta$ AICc | | 0.00 | 1.44 | | | |
| Akaike weight | | 0.63 | 0.31 | | | |
| Cumulative weight | | 0.63 | 0.93 | | | |

**Table S8.** Results from phylogenetic least squares regression model selection and model averaging for models relating EPA concentrations in edible finfish species tissues to finfish species' traits ($n$ = 168 species) (see Supplementary Methods 4 for full model description). Models are ranked in order of increasing AICc differences ($\Delta$ AICc). All models in the 95% confidence model set are shown here (cumulative sum of Akaike weights <= 0.95). Akaike weights indicate the relative likelihood of a model, given the set of models being considered.

Model-averaged standardized regression coefficients, $\beta_i$, with 95% CI, were averaged across all models in the best set and weighted by each model's Akaike weight. $\beta_i$ were calculated with $\beta_i$ = 0 when variable $i$ was not included in a model. $\beta_i$ with 95% confidence intervals not encompassing 0 are shown in bold. Relative variable importance ($w_{ip}$) is the sum of the relative variable importance across all models including that variable (*10*). Traits for which there is not a coefficient estimate did not appear in any of the models in the top 95% set (with cumulative Akaike weight <= 0.95). Note that different edible portions may come from different species, and not all tissues are available for all species.

**Table S9. Dependence of calcium (Ca) concentrations in edible tissues of small finfish on ecological traits.**

| Term | $w_{ip}$ | 1 | 2 | 3 | $\beta_i$ | Lower 95% CI | Upper 95% CI |
|---|---|---|---|---|---|---|---|
| Body part: muscle | 1.00 | + | + | + | 0.97 | -0.30 | 2.25 |
| Body part: muscle & organs (incl. bones) | 1.00 | + | + | + | 2.75 | 1.42 | 4.08 |
| Length | 0.67 | -0.29 | | -0.34 | -0.30 | -1.20 | 0.60 |
| Trophic position | 0.83 | 0.50 | 0.75 | | 0.60 | -0.52 | 1.73 |
| Feeding mode: hunting | 0.50 | | + | + | -0.12 | -1.18 | 0.95 |
| Feeding mode: variable | 0.50 | | + | + | 0.54 | -0.62 | 1.69 |
| Diet source | | | | | | | |
| Climate zone | | | | | | | |
| Realm | | | | | | | |
| $R^2$ | | 0.54 | 0.56 | 0.55 | | | |
| Δ AICc | | 0.00 | 0.84 | 2.09 | | | |
| Akaike weight | | 0.46 | 0.30 | 0.16 | | | |
| Cumulative weight | | 0.46 | 0.76 | 0.92 | | | |

**Table S9.** Results from ordinary least squares regression model selection and model averaging for models relating calcium concentrations in edible finfish species tissues to finfish species' traits (*n* = 39 species) for species with maximum length less than 50cm and mean length less than 25 cm (see Supplementary Methods 4 for full model description). Models are ranked in order of increasing AICc differences (Δ AICc). All models in the 95% confidence model set are shown here (cumulative sum of Akaike weights <= 0.95). Akaike weights indicate the relative likelihood of a model, given the set of models being considered. Model-averaged standardized regression coefficients, $\beta_i$, with 95% CI, were averaged across all models in the best set and weighted by each model's Akaike weight. $\beta_i$ were calculated with $\beta_i$ = 0 when variable *i* was not included in a model. $\beta_i$ with 95% confidence intervals not encompassing 0 are shown in bold. Relative variable importance ($w_{ip}$) is the sum of the relative variable importance across all models including that variable (*10*). Traits for which there is not a coefficient estimate did not appear in any of the models in the top 95% set (with cumulative Akaike weight <= 0.95). Note that different edible portions may come from different species, and not all tissues are available for all species.

**Table S10. Dependence of iron (Fe) concentrations in edible tissues of small finfish on ecological traits.**

| Term | $w_{ip}$ | 1 | 2 | 3 | 4 | 5 | $\beta_i$ | Lower 95% CI | Upper 95% CI |
|---|---|---|---|---|---|---|---|---|---|
| Trophic position | 0.46 | 0.66 | | | 0.47 | | 0.59 | -0.26 | 1.43 |
| Climate zone: polar | 1.00 | + | + | + | + | + | **-2.66** | **-4.34** | **-0.98** |
| Climate zone: subtropical | 1.00 | + | + | + | + | + | 0.24 | -1.35 | 1.84 |
| Climate zone: temperate | 1.00 | + | + | + | + | + | -1.30 | -2.63 | 0.03 |
| Climate zone: tropical | 1.00 | + | + | + | + | + | **-1.57** | **-2.87** | **-0.28** |
| Length | 0.53 | | -0.60 | | -0.75 | -0.53 | -0.64 | -1.40 | 0.12 |
| Feeding mode: hunting | 0.36 | | | + | + | | 0.71 | -0.11 | 1.53 |
| Feeding mode: variable | 0.36 | | | + | + | | **0.96** | **0.07** | **1.86** |
| Body part: muscle | 0.11 | | | | | + | -0.12 | -1.22 | 0.99 |
| Body part: muscle & organs (incl. bones) | 0.11 | | | | | + | 0.55 | -0.63 | 1.74 |
| Diet source | | | | | | | | | |
| realm | | | | | | | | | |
| $R^2$ | | 0.49 | 0.49 | 0.53 | 0.62 | 0.56 | | | |
| Δ AICc | | 0.00 | 0.26 | 0.85 | 0.94 | 1.85 | | | |
| Akaike weight | | 0.25 | 0.22 | 0.16 | 0.16 | 0.10 | | | |
| Cumulative weight | | 0.25 | 0.47 | 0.63 | 0.79 | 0.89 | | | |

**Table S10.** Results from ordinary least squares regression model selection and model averaging for models relating iron concentrations in edible finfish species tissues to finfish species' traits ($n$ = 35 species) for species with maximum length less than 50cm and mean length less than 25 cm (see Supplementary Methods 4 for full model description). Models are ranked in order of increasing AICc differences (Δ AICc). All models in the 95% confidence model set are shown here (cumulative sum of Akaike weights <= 0.95). Akaike weights indicate the relative likelihood of a model, given the set of models being considered. Model-averaged standardized regression coefficients, $\beta_i$, with 95% CI, were averaged across all models in the best set and weighted by each model's Akaike weight. $\beta_i$ were calculated with $\beta_i = 0$ when variable $i$ was not included in a model. $\beta_i$ with 95% confidence intervals not encompassing 0 are shown in bold. Relative variable importance ($w_{ip}$) is the sum of the relative variable importance across all models including that variable (*10*). Traits for which there is not a coefficient estimate did not appear in any of the models in the top 95% set (with cumulative Akaike weight <= 0.95). Note that different edible portions may come from different species, and not all tissues are available for all species.

**Table S11. Dependence of zinc (Zn) concentrations in edible tissues of small finfish on ecological traits.**

| Term | $w_{ip}$ | 1 | 2 | 3 | 4 | 5 | 6 | $\beta_i$ | Lower 95% CI | Upper 95% CI |
|---|---|---|---|---|---|---|---|---|---|---|
| Length | 0.47 | -0.36 | | | -0.37 | | | -0.36 | -0.93 | 0.21 |
| Feeding mode: hunting | 1 | + | + | + | + | + | + | 0.3 | -0.35 | 0.94 |
| Feeding mode: variable | 1 | + | + | + | + | + | + | **0.93** | **0.25** | **1.61** |
| Trophic position | 0.44 | | 0.38 | | | 0.38 | 0.4 | 0.38 | -0.28 | 1.03 |
| Diet source: demersal | 0.12 | | | + | | | + | -0.22 | -0.82 | 0.38 |
| Diet source: pelagic-neritic | 0.12 | | | + | | | + | 0.26 | -0.44 | 0.97 |
| Diet source: pelagic-oceanic | 0.12 | | | + | | | + | 0.53 | -0.24 | 1.31 |
| Body part: muscle & organs (incl. bones) | 0.15 | | | | + | + | | -0.04 | -0.54 | 0.46 |
| Climate zone | | | | | | | | | | |
| Realm | | | | | | | | | | |
| $R^2$ | | 0.33 | 0.32 | 0.41 | 0.33 | 0.32 | 0.46 | | | |
| Δ AIC | | 0 | 0.31 | 3.07 | 3.21 | 3.57 | 4.85 | | | |
| Akaike weight | | 0.36 | 0.31 | 0.08 | 0.07 | 0.06 | 0.03 | | | |
| Cumulative weight | | 0.36 | 0.68 | 0.75 | 0.83 | 0.89 | 0.92 | | | |

**Table S11**. Results from ordinary least squares regression model selection and model averaging for models relating zinc concentrations in edible finfish species tissues to finfish species' traits (*n* = 28 species) for species with maximum length less than 50cm and mean length less than 25 cm (see Supplementary Methods 4 for full model description). Models are ranked in order of increasing AICc differences (Δ AICc). All models in the 95% confidence model set are shown here (cumulative sum of Akaike weights <= 0.95). Akaike weights indicate the relative likelihood of a model, given the set of models being considered. Model-averaged standardized regression coefficients, $\beta_i$, with 95% CI, were averaged across all models in the best set and weighted by each model's Akaike weight. $\beta_i$ were calculated with $\beta_i$ = 0 when variable *i* was not included in a model. $\beta_i$ with 95% confidence intervals not encompassing 0 are shown in bold. Relative variable importance ($w_{ip}$) is the sum of the relative variable importance across all models including that variable (*10*). Traits for which there is not a coefficient estimate did not appear in any of the models in the top 95% set (with cumulative Akaike weight <= 0.95). Note that different edible portions may come from different species, and not all tissues are available for all species.

**Table S12. Seafood consumed by North America Indigenous communities.**

| Culture | Abbreviation | Region | Location | Taxa |
|---|---|---|---|---|
| Abenaki | AB | Northeast | Quebec, Maine | 29 |
| Bella Coola | BC | Northwest Coast | British Columbia | 40 |
| Central Salish | CS | Northwest Coast | British Columbia, Washington | 42 |
| Cree | CR | Subarctic | Labrador, Quebec, Ontario, Manitoba, Saskatchewan, Alberta | 27 |
| Haida | HC | Northwest Coast | British Columbia, Alaska | 36 |
| Inuit-Inupiaq | II | Arctic | Alaska, Northwest Territories, Nunavut, Nunavik, Quebec, Labrador | 52 |
| Kwakiutl | KW | Northwest Coast | British Columbia | 40 |
| Miˈkmaq | MI | Northeast | Nova Scotia, New Brunswick, Quebec, Newfoundland | 57 |
| Montagnais-Naskapi | MN | Subarctic | Labrador, Quebec | 25 |
| Nootkan | NO | Northwest Coast | British Columbia, Washington | 49 |
| Tlingit | TL | Northwest Coast | British Columbia, Yukon, Alaska | 51 |
| Tsimshian | TS | Northwest Coast | British Columbia, Alaska | 41 |
| Wampanoag | WA | Northeast | Massachusetts | 35 |
| Yupik | YU | Arctic | Alaska | 38 |

**Table S12.** North American Indigenous cultures, and numbers of seafood species in traditional diets used in the local scale nutritional benefits analysis (**Figure S4**; Supplementary Methods 1.4 and 3).

**Table S13. Top fourteen of 41 most commonly consumed species as per FAO production volumes in the nutrient dataset. Data from (9).**

| Genus species |
|---|
| *Theragra chalcogramma* |
| *Gadus morhua* |
| *Gadus macrocephalus* |
| *Tenualosa ilisha* |
| *Rastrelliger kanagurta* |
| *Merluccius productus* |
| *Oncorhynchus gorbuscha* |
| *Pollachius virens* |
| *Melanogrammus aeglefinus* |
| *Thunnus alalunga* |
| *Oreochromis niloticus* |
| *Panaeus monodon* |
| *Portunus pelagicus* |
| *Trachurus trachurus* |

**Table S14. Predictors used in phylogenetic generalized least squares models used to predict nutrient content as a function of ecological traits.**

| *Predictor* | *Categories* |
|---|---|
| *Feeding mode* | Filtering plankton, grazing, hunting, selective plankton feeding, variable |
| *Diet source and habitat* | Reef associated, Pelagic, Oceanic, Neritic, Demersal, Benthopelagic, Bathypelagic |
| *Realm* | Freshwater, Freshwater or brackish, Freshwater or brackish or marine, marine |
| *Climate zone* | Temperate, subtropical, polar, tropical, boreal |
| *Maximum body length* | continuous |
| *Trophic position* | continuous |
| *Body part* | Muscle, muscle + organs (including bones, skin etc.), eggs or liver |

#### Appendix A: References

1. Institute of Medicine, *Dietary Reference Intakes for Calcium and Vitamin D. Washington* (2011) https:/doi.org/10.1016/B978-1-893997-82-0.50009-8.
2. JECFA, Evaluation of certain contaminants in food: seventy-second [72nd] report of the Joint FAO/WHO Expert Committee on Food Additives. *World Heal. Organ. https//apps.who.int/iris/bitstream/handle/10665/44514/WHO_TRS_959_eng.pdf,* (accessed 03.04.20) (2011).
3. M. W. Cadotte, K. Carscadden, N. Mirotchnick, Beyond species: Functional diversity and the maintenance of ecological processes and services. *J. Appl. Ecol.* **48**, 1079–1087 (2011).
4. E. Laliberte, P. Legendre, A distance-based framework for measuring functional diversity from multiple traits. *Ecology* **91**, 299–305 (2010).
5. O. L. Petchey, A. Hector, K. J. Gaston, How do different measures of functional diversity perform? *Ecology* **85**, 847–857 (2004).
6. R. Remans, *et al.*, Assessing nutritional diversity of cropping systems in African villages. *PLoS One* **6** (2011).
7. Institute of Medecine, *Dietary Reference Intakes for Energy, Carbohydrate, Fiber, Fat, Fatty Acids, Cholesterol, Protein, and Amino Acids* (National Academies Press, 2005) https:/doi.org/10.17226/10490 (October 8, 2019).
8. Institute of Medecine, *Dietary Reference Intakes for Vitamin A, Vitamin K, Arsenic,*

*Boron, Chromium, Copper, Iodine, Iron, Manganese, Molybdenum, Nickel, Silicon, Vanadium, and Zinc* (National Academies Press, 2001) https:/doi.org/10.17226/10026 (October 8, 2019).
9. Food and Agriculture Organization, "Capture production by principal species in 2013" (2013) (May 22, 2018).
10. Burnham KP, Anderson DR (2002) Model selection and multimodel inference: a practical information-theoretic approach, 2nd edn. Springer, New York

### Part B: Supplementary Methods

**This file includes:**

Supplementary Methods
Part B: References

**Other supplementary materials for this manuscript include the following:**

**Part A:** Supplementary Figures and Tables

#### Supplementary Methods

#### 1. Literature search and data collection

##### 1.1 Nutrient concentration data

We assembled a dataset of published nutrient concentrations in edible portions of 547 aquatic species (**Figure S1A**). We aimed to include as many marine and freshwater species as possible covering a wide geographic extent. We searched peer-reviewed literature for analytical food composition values as well as the Food and Agriculture Organization's Global Food Composition Database for Fish and Shellfish (INFOODS) (1). We extracted data from peer-reviewed literature by downloading datasets directly when available, or extracting data from tables using the 'Tabula' software, from figures using 'WebPlotDigitizer', or manually when none of these options was available. We searched the peer-reviewed literature using Google Scholar for papers published between 1970 and 2017, using search terms including (but not limited to): "seafood AND micronutrient", or "fish AND zinc", "fish AND calcium", "fish AND iron", or "fish AND essential fatty acids", "fish AND nutrient composition" or species-specific searches such as "*Mya arenaria* AND zinc" to find published papers on edible tissue nutrient concentrations that met the following criteria. First, for finfish, we restricted our search and analysis to include only edible portions of wild caught, raw fish (excluding prepared or farmed finfish samples). We included both farmed and wild caught mollusc species because mollusc farming does not typically involve additional food inputs, which could influence tissue nutrient composition. Second, we only included data for which the body part contained in the edible portion was made explicit (i.e. muscle tissue, whole body, eggs). For each sample, we noted which body parts are included in the edible portion and season of collection. We did not include data from national food composition tables because these data usually report seafood data with a generic food description (for example 'salmon'), which does not allow for a clear description of which fish tissues are included in the edible portion or the species. We excluded samples that were of mixed species groups, or not identified to genus and species. We only included data that were available in terms of concentrations per fresh wet weight. We standardized all measurements to mg (for calcium, iron and zinc) or g (for DHA, EPA, fat and protein) per 100g edible tissue. To address inconsistencies in fatty acid data reporting, consistent with fatty acid data in the INFOODS database, we standardized fatty acid measurements using the fatty acid conversion factors proposed by Nowak et al. 2014 (2). When there were multiple observations available for a single species and given body part, we averaged nutrient concentrations across the observations. For each sample, we noted the location of collection (e.g. latitude and longitude). We standardized species' Latin names (i.e. genus and species) using the *taxize* package in R (3). In total, we assembled 5288 observations of nutrient concentrations.

##### 1.2 Ecological trait data

To test our hypothesis that nutritional benefits may depend on biodiversity because they are correlated with ecological functional trait diversity among aquatic species, we collected ecological trait information from FishBase (4) and SeaLifeBase (5) (**Figure S1E**). We selected a range of ecological functional traits that relate to species' ecological roles (6): body size (maximum length), fractional trophic position, habitat and diet source (e.g. oceanic, reef-associated, neritic) and feeding mode (e.g. filter-feeding vs hunting), realm (freshwater, brackish, marine), and climate zone (e.g. tropical, temperate) (Table S14). We focused on this set of functional traits because these traits are related to species' diets, morphology, energetic demands and habitats, all of which may influence the concentration of biologically essential elements and fatty acids in their tissues (7, 8), and which determine the functional roles that organisms play in ecosystems (6). We extracted these data directly from FishBase or SeaLifeBase data using the R package *rfishbase (9)*. If multiple observations were available for a single ecological functional trait for a single species, we took an average across multiple observations. This approach allowed us to characterize traits at the species' level, although it does not allow investigation of intraspecific variation in traits across different regions or times.

##### 1.3 Contaminant data from North American seafood species

We drew on existing, published datasets for heavy metals that can be toxic to human health if they are consumed in high enough quantities (contaminants). We synthesized contaminant concentrations in North American finfish and invertebrate species including mercury, cadmium, arsenic and lead (10, 11) (**Figure S1A**). These datasets contain data from the peer-reviewed literature, as well as US federal and state government reports. From these datasets, we extracted data for samples which were identifiable to species, and which consisted of raw, wild muscle tissue only (thus excluding farmed, cooked, tinned or otherwise prepared samples), and which were wild-caught in North America (i.e. thus excluding samples that were sourced from markets and for location of origin may be unknown). In total, we extracted 2209 observations of contaminant concentrations: 322 observations for each of lead, arsenic and cadmium, and 1243 observations for mercury, which resulted in mercury concentrations for 353 species, and lead, arsenic and cadmium concentrations for 200 species. These observations were of total metal concentrations, and did not account for metal fractions in different forms. For mercury, because approximately 95% of total mercury burden in finfish is methylmercury (12, 13), and methylmercury is the form of mercury that poses most risk to human health, we calculated methylmercury concentrations assuming that 95% of total mercury is methylmercury (following (10, 13, 14)). However, because molluscs tend to have much lower methylmercury fractions, we assumed methylmercury was 35% of total mercury (based on (14–16)). For arsenic, we considered total arsenic, and did not address ratios between total and inorganic forms of arsenic. A source of error in this analysis is the percentage of total mercury that is methylmercury, which may vary across species (15, 17). Because we still do not know what drives variation in methylmercury to total mercury ratios across species, and methylmercury data are not widely available, this was a necessary assumption in this analysis, and deserves further study.

**1.4 Biodiversity in North American Indigenous diets**

To assess the effects of biodiversity in seafood diets sourced from local or regional species pools, we determined species lists for species included in traditional animal food diets (past or present) in North America. We focused on traditional Indigenous diets in North America because many of these communities rely on locally harvested aquatic species to meet their nutritional needs (18–20). We compiled lists of seafood species contained in traditional diets from an ethnographic database of traditional animal foods of Indigenous peoples of North America (21). This database represents a compilation of hundreds of ethnographic sources, and includes data collected with a range of methods, including interviews, 24-hour recall surveys, food questionnaires, and archaeological records. From this database, we extracted information on seafood species associated with Indigenous cultures' traditional diets, how these species are traditionally prepared (e.g. smoked, raw, steamed), and which body parts are included in the edible portion (e.g. eggs, muscle, liver). The information available was not quantitative estimates of amounts or frequencies of foods consumed, rather just the lists of species traditionally included in diets (currently or historically), which allowed us to quantify the role of biodiversity theoretically. From this dataset of more than 70 Indigenous diets, we focused on Indigenous cultures which included more than 25 seafood species in their traditional diet. This resulted in a dataset of the seafood species included in the traditional diets of fourteen Indigenous cultures, including Abenaki, Bella Coola, Central Salish, Cree, Haida, Inuit-Inupiaq, Kwakiutl, Miˈkmaq, Montagnais-Naskapi, Nootkan, Tlingit, Tsimshian, Wampanoag and Yupik (Table S12).

**2. Quantifying the effects of seafood biodiversity on contaminant exposure**

We quantified effects of biodiversity on contaminant exposure using methods analogous to those described for nutritional benefits, using two metrics: *NC* and $P_{max}$. Contaminant exposure risks were estimated for species sampled from the North American seafood contaminant dataset (Supplementary Methods 1.3) (**Figure S1C**).

1) Number of contaminants, *NC* (Table S1, Equation A4): We quantified the effects of seafood species richness on contaminant exposure using methods analogous to those we used for the nutrients. We assembled diets from the North American species pool (Supplementary Methods 1.3) at random, with a replacement design, as described in Methods 2.2. To test the hypothesis that complementarity in contaminant concentrations among species increases health risks by increasing the number of distinct Provisional Tolerable Daily Intake (PTDI; Table S1) limits exceeded in a 100g portion, we quantified, for all possible combinations of species at each level of species richness (from 1 - 10 species, $n$ = 1023), the number of distinct contaminant PTDIs exceeded by assigning each combination of species a set of 0's or 1's according to whether that

combination exceeded the PTDI for each contaminant (see Table S1, Equation A4) (**Figure S1B, C**). Our approach allowed us to explore how likely it would be for human diets containing different numbers of fish species to exceed a given number of contaminant PTDI limits (*NC* ranges between 0 and 4), assuming that fish species were included in the human diet at random. At level of species richness, we took an average of *NC*, and then repeated this process of assembling diets, and estimating *NC* 1000 times. This process yielded 1000 estimates of *NC* (**Figure S1B, C**). We quantified the effect of biodiversity on *NC* ($n$ = 1000 estimates of *NC* per richness level) by fitting a power function,

$$NC = aS^{b_{NC}} \qquad (B1)$$

where the parameter $b_{NC}$ describes the relationship between a change in species richness, *S*, and a change in *NC* (i.e. the number of distinct PTDI limits exceeded per average 100g portion), and *a* is a constant (**Figure S1D**).

2) Maximum portion size, $P_{max.}$(Table S1 Equation A3): We assessed the effect of increasing seafood species richness on the average contaminant concentrations in seafood diets, and therefore the maximum portion size before upper tolerable limits are exceeded. Following similar methods as described above for minimum portion size, $P_{min}$ (Methods 1.1): from the North American species pool (Supplementary Methods 1.3), we sampled ten species at random and then assembled seafood diets from all possible combinations of these ten randomly chosen species at 10 levels of species richness (1-10). For each combination of species at each level of species richness (1-10 species), we calculated the number of grams required to exceed a given contaminant tolerable upper intake (100% of PTDI), $P_{max}$ (see Table S1 Equation A3). At each level of species richness, we took an average $P_{max}$. We repeated this process of sampling ten

species from the North American species pool, assembling all possible diets at each richness level, and estimating $P_{max}$ 1000 times, yielding 1000 estimates of $P_{max}$ per richness level. We quantified the effect of species richness in a diet on maximum portion size, $P_{max}$ ($n$ = 1000 estimates of $P_{max}$ per richness level), by fitting a power function to these $P_{max}$ estimates:

$$P_{max} = aS^{b_{Pmax}} \qquad (B2)$$

where the parameter $b_{Pmax}$ describes the relationship between a change in species richness, $S$, and a change in $P_{max}$, and $a$ is a constant (in units of grams) (**Figure S1D**). Since $P_{max}$ is measured in grams required to exceed a given PTDI threshold, and fewer grams required is worse from the perspective of human nutrition, then a negative effect of biodiversity would be reflected in a negative $b_{Pmax}$ (i.e. $P_{max}$, measured in grams of tissue required, decreases with species richness).

**3. Comparing local scale biodiversity effects to global scale biodiversity effects**

We assessed biodiversity effects on nutritional benefits in local scale Indigenous diets as described in Methods 2.2 and 2.3 for global seafood diets. Instead of sampling from the global species pool, we sampled diets from species contained within traditional diets in fourteen Indigenous cultures in North America (Supplementary Methods 1.4, Table S12). For each of these diets, we repeated the replacement design randomization process described above, and calculated *NT* and $P_{min}$ (Methods 2.2). We compared estimates of the 'biodiversity effect', the slope parameter, $b$, at global and local scales, by comparing the $b$ estimates from the 14 traditional Indigenous diets to global diets standardized to 40 species (the average number of species in the Indigenous diets, Methods 2.2), indicated as 'GL' in **Figure S4**.

We tested the hypothesis that biodiversity effects at local and global scales are associated with differences in the diversity of nutritional profiles of species available at local and global scales. We used a metric called 'nutritional functional diversity', *NFD* (22, 23). *NFD* is based on an assessment of the entire functional diversity of a group represented as a functional dendrogram, and *NFD* allows estimation of complementarity among species' nutrient concentrations (i.e. nutritional functional traits) using the dendrogram. We hypothesized that *NFD* would be higher at the global scale than the local scale, because the global species pool contains more ecological and biogeographic diversity. We treated the concentration of each micronutrient (calcium, iron, zinc, EPA and DHA) as a nutritional functional trait. We also quantified a metric of nutritional functional evenness metric (*NFEve*) using the *dbFD* function in the *FD* package in R (24), which normally quantifies the evenness of abundance in a functional trait space. Here, we used *NFEve* to quantify the evenness in concentration of nutrients across species (25). To compare *NFD* and *NFEve* at the global and local scales, we first subsampled 40 species (the average species pool at the local scale) from the global pool, then calculated the functional diversity metrics on the subsample, and repeated this process 1000 times. Using this same approach, we calculated levels of 'expected' *NFD* and *NFEve* for each local diet by choosing random subsets of the global pool with sample size equal to the species pool in each local diet, and repeated this process 1000 times (**Figure S5**).

**4. Assessing the relationship between species' ecological traits and nutrient concentrations**

We tested for complementarity in nutrient concentrations among species by calculating Pearson correlation coefficients among nutrient concentrations across species (i.e. pairwise correlations across all species). We identified correlations between multiple nutrients in seafood tissues using principal components analysis, using the '*rda*' function in *vegan* (94). We tested the hypothesis that major phylogenetic groups correlated with functional differences in life history, resource use and ecology (i.e. finfish, mollusc, and crustacean) differ in their multi-nutrient profiles via permutational multivariate ANOVA (PERMANOVA) using the *adonis* function (999 permutations) based on Bray-Curtis dissimilarity matrices.

To test for associations between species' ecological functional traits and their nutrient concentrations, we modeled the relationship between traits and ln(nutrient concentration) with phylogenetic least squares regression (PGLS). To assess whether the relationships between species' traits and their nutrient concentrations were associated, we fit multiple regression models using PGLS using the *gls* function in the *nlme* package in R (26). Unlike in ordinary least squares (OLS), which assumes there is no covariance structure in the error term, ε, (all species are independent from one another, and residuals from closely related species are not more similar on average than residuals from distantly related species), PGLS assumes that the residuals are non-independent, and that the expected covariance is related to the shared evolutionary history between the species.

The full model included the entire set of trait predictors (Table S14) as fixed effects:

$$ln(concentration) = \beta_0 + \beta_1 \times ln(body\ length) + \beta_2 \times trophic\ position + \beta_3 \times feeding\ mode + \beta_4 \times diet\ source + \beta_5 \times realm + \beta_6 \times climate\ zone + \beta_7 \times body\ part + \varepsilon$$

This approach allowed us to account for phylogenetic non-independence by using shared ancestry as weights on the elements of the residual variance-covariance matrix used in the model. We created a supertree by combining phylogenies that included the species of finfish, molluscs and crustaceans in our nutrient dataset using the *rotl* package in R (27), which is an interface to the Open Tree of Life (28). We computed branch lengths according to taxonomic depth (29) using the *compute.brlen* function in the *ape* package in R (30). We incorporated phylogenetic information into the models using Pagel's λ correlation structure (31) constructed with the *corPagel* function in the *ape* package and estimated the amount of phylogenetic signal in the data estimated using maximum likelihood. When $\lambda = 0$, this suggests that the relationship between predictor and response variables is unrelated to phylogeny, and when $\lambda = 1$, this indicates that traits have evolved under Brownian motion on the given phylogeny such that variation among species in traits may reflect evolutionary history rather than contemporary predictor variables. Where λ was negative, suggesting closely related species have negatively correlated phenotypes under the Brownian model of evolution, we fixed λ at 0. To assess whether the relationships between species' traits and their nutrient concentrations were associated, we fit multiple regression models using PGLS using the *gls* function in the *nlme* package in R (26).

We created models from subsets of the full model that represented hypotheses based on the known physiological roles of micronutrients and their relationships to our set of predictors. To avoid issues associated with multicollinearity of predictor variables, we excluded other possible variables if they were highly correlated (i.e. correlation coefficient > 0.6). We identified the best subset of models using the Akaike Information Criterion, adjusted for small sample sizes (AICc).

We used AICc, Δ AICc and Akaike weights (*w*) to compare models. We ranked models based on AICc, and selected the set of models that produced a cumulative *w* >= 0.95, meaning that we are 95% confident that the chosen set includes the best model (32). We limited our analyses to species for which we had complete trait data (Table S14), identified to the species level; we did not impute missing trait data using phylogenetic or taxonomic relationships. To account for model uncertainty in the ecological trait correlation analyses, we performed model averaging of coefficients in all models in the 95% confidence set, and included zeros as coefficients when variables did not enter a given model (32). We conducted our model selection and averaging analyses with the *MuMIn* package (33) and all other analyses in R version 3.3.2 (*18*).

The nutrient concentration of different seafood tissues varies according to which body parts are included in the edible portion (e.g. muscle tissues, eggs, liver). To control for these effects of body part in our analyses of ecological traits, we included 'body part' as a covariate, and in a subset of analyses, we included data for muscle tissues only. We also tested for associations between nutrient concentrations (calcium, iron and zinc) and ecological traits among small finfish species (mean length < 25 cm; maximum length < 50 cm), whose edible portions more often include tissues other than muscle only, using ordinary least squares regression (noting that the strength of the phylogenetic signal for calcium, iron and zinc, λ = 0).

**5. Quantifying the relationship between ecological functional diversity and nutritional benefits**

To quantify the relationship between ecological functional diversity and nutritional benefits, we created simulated diets from the global species as described above (Methods 2.2). To isolate the effects of ecological functional diversity from species richness, we focus on diets of ten species only, thereby capturing the effect of changes in ecological functional diversity independent from changes in species richness. For each of these diets, we quantified the ecological functional diversity, minimum portion size required, $P_{min}$, and the number of nutrient RDA targets per 100g, *NT* (Methods 1.3). We estimated ecological functional diversity using a metric of trait dispersion in multidimensional ecological trait space: functional dispersion (34) (**Figure S1F**). Hereafter, when we refer to ecological functional diversity, we mean functional dispersion. This metric measures the distance of each species from the mean coordinates of the assemblage, weighted by abundance. However, in our simulated diets, since abundance of all species was equal, this metric is simply the mean distance, in multidimensional trait space, of each individual species from the centroid of all species. Here, we constructed the multidimensional trait space using a set of the ecological functional traits (body size, trophic position, diet source and feeding mode). We chose to focus on these traits because they are strongly related to the ecological roles that species play in ecological communities. We estimated the relationship between ecological functional diversity and *NT* (which ranges from 0-5) using ordinal logistic regression. We estimated the relationship between ecological functional diversity and minimum portion size, $P_{min}$, (in grams) using reduced major axis regression.

**Part B: References**

1. Food and Agriculture Organization of the United Nations, "FAO/INFOODS Global Food Composition Database for Fish and Shellfish Version 1.0- uFiSh1.0" (2016).

2. V. Nowak, D. Rittenschober, J. Exler, U. R. Charrondiere, Proposal on the usage of conversion factors for fatty acids in fish and shellfish. *Food Chem.* **153**, 457–463 (2014).

3. S. Chamberlain, *et al.*, taxize: Taxonomic information from around the web. *R package version 0. 3. 0* (2014).

4. Froese, R. and D. Pauly. Fishbase. World Wide Web Electronic Publication (2020), version (12/2020), www.fishbase.org

5. Palomares, M.L.D. and D. Pauly. SeaLifeBase. World Wide Web Electronic Publication, version (12/2020).

6. C. Violle, *et al.*, Let the concept of trait be functional! *Oikos* **116**, 882–892 (2007).

7. R. Karimi, C. L. Folt, Beyond macronutrients: element variability and multielement stoichiometry in freshwater invertebrates. *Ecol. Lett.* **9**, 1273–1283 (2006).

8. M. T. Arts, M. T. Brett, M. Kainz, *Lipids in Aquatic Ecosystems* (Springer Science & Business Media, 2009).

9. C. Boettiger, D. T. Lang, P. C. Wainwright, rfishbase: exploring, manipulating and visualizing FishBase data from R. *J. Fish Biol.* **81**, 2030–2039 (2012).

10. R. Karimi, T. P. Fitzgerald, N. S. Fisher, A quantitative synthesis of mercury in commercial seafood and implications for exposure in the United States. *Environ. Health Perspect.* **120**, 1512–1519 (2012).

11. R. A. Hall, E. G. Zook, G. M. Meaburn, National Marine Fisheries Service survey of trace elements in the fishery resource (1978).

12. M. N. Piraino, D. L. Taylor, Bioaccumulation and trophic transfer of mercury in striped bass (Morone saxatilis) and tautog (Tautoga onitis) from the Narragansett Bay (Rhode Island, USA). *Mar. Environ. Res.* **67**, 117–128 (2009).

13. N. S. Bloom, On the Chemical Form of Mercury in Edible Fish and Marine Invertebrate Tissue. *Can. J. Fish. Aquat. Sci.* **49**, 1010–1017 (1992).

14. E. M. Sunderland, M. Li, K. Bullard, Decadal Changes in the Edible Supply of Seafood and Methylmercury Exposure in the United States. *Environ. Health Perspect.* **126**, 017006 (2018).

15. C. Y. Chen, *et al.*, Mercury bioavailability and bioaccumulation in estuarine food webs in the Gulf of Maine. *Environ. Sci. Technol.* **43**, 1804–1810 (2009).

16. M. Li, *et al.*, Environmental Origins of Methylmercury Accumulated in Subarctic Estuarine Fish Indicated by Mercury Stable Isotopes. *Environ. Sci. Technol.* **50**, 11559–11568 (2016).

17. Y. Yamashita, Y. Omura, E. Okazaki, Total mercury and methylmercury levels in commercially important fishes in Japan. *Fish. Sci.* **71**, 1029–1035 (2005).

18. H. V. Kuhnlein, O. Receveur, Local Cultural Animal Food Contributes High Levels of Nutrients for Arctic Canadian Indigenous Adults and Children. *J. Nutr.* **137**, 1110–1114 (2007).

19. H. V. Kuhnlein, B. Erasmus, D. Spigelski, *Indigenous Peoples' food systems: the many dimensions of culture, diversity and environment for nutrition and health*, FAO, Ed. (2009).

20. A. M. Cisneros-Montemayor, D. Pauly, L. V. Weatherdon, Y. Ota, A Global Estimate of Seafood Consumption by Coastal Indigenous Peoples. *PLoS One* **11**, e0166681 (2016).

21. H. V. Kuhnlein, M. M. Humphries, *Traditional Animal Foods of Indigenous Peoples of Northern North America*, Centre for Indigenous Peoples' Nutrition and Environment, McGill University, Ed. (2017).

22. O. L. Petchey, K. J. Gaston, Functional diversity (FD), species richness and community composition. *Ecol. Lett.* **5**, 402–411 (2002).

23. R. Remans, *et al.*, Assessing nutritional diversity of cropping systems in African villages. *PLoS One* **6** (2011).

24. A. E. Laliberté, P. Legendre, B. Shipley, M. E. Laliberté, FD: measuring functional diversity from multiple traits, and other tools for functional ecology. R package: 0--12 (2014).

25. S. Villéger, N. W. H. Mason, D. Mouillot, New multidimensional functional diversity indices for a multifaceted framework in functional ecology. *Ecology* **89**, 2290–2301 (2008).

26. J. Pinheiro, D. Bates, S. DebRoy, D. Sarkar, R. C. Team, nlme: linear and nonlinear mixed effects models. R package version 3.1-117. *R Core Team*, 1–117 (2014).

27. F. Michonneau, J. W. Brown, D. J. Winter, rotl : an R package to interact with the Open Tree of Life data. *Methods Ecol. Evol.* **7**, 1476–1481 (2016).

28. C. E. Hinchliff, *et al.*, Synthesis of phylogeny and taxonomy into a comprehensive tree of life. *Proc. Natl. Acad. Sci. U. S. A.* **112**, 12764–12769 (2015).

29. A. Grafen, The phylogenetic regression. *Philos. Trans. R. Soc. Lond. B Biol. Sci.* **326**, 119–157 (1989).

30. E. Paradis, J. Claude, K. Strimmer, APE: Analyses of Phylogenetics and Evolution in R language. *Bioinformatics* **20**, 289–290 (2004).

31. L. J. Revell, Phylogenetic signal and linear regression on species data. *Methods Ecol. Evol.* **1**, 319–329 (2010).

32. Burnham KP, Anderson DR (2002) Model selection and multimodel inference: a practical information-theoretic approach, 2nd edn. Springer, New York.

33. Barton, K., (2018). MuMIn: Multi-Model Inference. R package version 1.42.1. https://CRAN.Rproject.org/package=MuMIn

34. E. Laliberté, P. Legendre, A distance-based framework for measuring functional diversity from multiple traits. *Ecology* **91**, 299–305 (2010).
